# Multivalent polymers can control phase boundary, dynamics, and organization of liquid-liquid phase separation

**DOI:** 10.1101/2020.12.31.424934

**Authors:** Emiko Zumbro, Alfredo Alexander-Katz

## Abstract

Multivalent polymers are a key structural component of many biocondensates. When interacting with their cognate binding proteins, multivalent polymers such as RNA and modular proteins have been shown to influence the liquid-liquid phase separation (LLPS) boundary to control condensate formation and to influence condensate dynamics after phase separation. Much is still unknown about the function and formation of these condensed droplets, but changes in their dynamics or phase separation are associated with neurodegenerative diseases such as amyotrophic lateral sclerosis (ALS) and Alzheimer’s Disease. Therefore, investigation into how the structure of multivalent polymers relates to changes in biocondensate formation and maturation is essential to understanding and treating these diseases. Here, we use a coarse-grain, Brownian Dynamics simulation with reactive binding that mimics specific interactions in order to investigate the difference between non-specific and specific multivalent binding polymers. We show that non-specific binding interactions can lead to much larger changes in droplet formation at lower energies than their specific, valence-limited counterparts. We also demonstrate the effects of solvent conditions and polymer length on phase separation, and we present how modulating binding energy to the polymer can change the organization of a droplet in a three component system of polymer, binding protein, and solvent. Finally, we compare the effects of surface tension and polymer binding on the condensed phase dynamics, where we show that both lower protein solubilities and higher attraction/affinity of the protein to the polymer result in slower droplet dynamics. We hope this research helps to better understand experimental systems and provides additional insight into how multivalent polymers can control LLPS.

## Introduction

Multivalency is employed throughout biology for numerous reasons including building conformal interfaces, increasing specificity of bonds using a limited number of ligand types, and creating much stronger bonds by using many low affinity bonds simultaneously [1]. Multivalent binding is defined as when multiple binding sites on both interacting species bind simultaneously to create a much stronger bond than the sum of the constituent monovalent binding affinities [1]. Multivalent species can come in many architectures, but here, we focus on multivalent polymers and their role in biocondensates or membraneless organelles. Multivalent proteins and nucleic acids have been found in many membraneless organelles, which are membraneless, liquid like droplets found inside cells which concentrate and organize specific sets of molecules [2]. Although these biocondensates can have tens to hundreds of components, studies have shown that multivalent polymers are key directors of the phase separation of condensates and multivalent polymers can undergo phase separation with just their target binding species in vivo, in vitro, and in simulation [2–6]. These studies suggests that controlling the features of multivalent polymers can modulate the formation of biocondensates and the recruitment of other important components after the initial phase separation [6–11].

Because aberrant phase separation of these biocondensates is associated with neurodegenerative diseases such as amyotrophic lateral sclerosis (ALS), Parkinson’s, and Alzheimer’s disease, understanding their formation is an important area of research [2, 4, 12–14]. Changes in dynamics of globules and a liquid-to-solid transition in biocondensates has also been associated with disease [15]. Since multivalent polymers control these biocondensates, exploring how multivalent polymer properties can change both the kinetics and thermodynamics of liquid-liquid phase separation (LLPS) is essential. Theoretical and experimental studies of these systems have shown that increasing valency and individual binding site affinities can lower the phase separation boundary to more dilute species concentrations [3, 5–7]. Another theoretical study found that the solvation of polymeric linkers between binding sites controls whether polymers form a cross-linked gel with or without phase separation [6]. In previous work, we showed that polymer flexibility or persistence length can change the effective valency and consequently, the phase boundary of multivalent polymers [16]. In this paper, we further build on the understanding that multivalent polymer characteristics can control the thermodynamics of biocondensates by exploring the effect of non-specific versus specific polymer binding interactions, condensed phase nucleation in smaller systems, and the dynamics of the resulting condensed polymer droplets.

We use a coarse-grain Brownian Dynamics simulation to explore the phase separation of long, many-valent polymers and smaller binding targets. This system represents a polymer binding to a smaller, mismatched-valence target such as RNA interacting with RNA-binding proteins in ribonucleicprotein (RNP) granules [17, 18]. Using Brownian Dynamics allows us to capture both the thermodynamics and kinetics of phase separated polymer-target systems, providing insight on how polymers can change both the formation and dynamic properties of biocondensates. In this work, we show that nucleating a condensed phase using non-specific interactions, such as hydrophobicity or charge, occurs at lower attraction energies than using valence-limited lock-and-key type binding such as those between a ligand and a folded protein pocket. Therefore, non-specific and specific interactions can be combined to carefully adjust phase transition boundaries. Next, by looking at the internal arrangement of the resulting condensates, we explain how changing polymer interactions can control the spatial organization of the condensed phase. Last, we investigate how polymer properties can alter the kinetics inside condensed droplets.

## Materials and methods

To study the condensed phase nucleation of multivalent polymers, we use coarse-grain Brownian dynamics simulations with a bead-spring polymer and a spherical binding target represented as a single bead of the same size as the polymer beads [19]. This scenario represents a general model of the protein-protein or nucleic acid-protein binding found in the formation of membraneless organelles. It most closely resembles a piece of RNA binding to an RNA binding protein such as hnRNPA1 found in stress granules, and whose solidification of the condensed phase is associated with ALS and fronto-temporal dementia [17, 18, 20]. We applied Brownian dynamics to each bead governed by the equation:

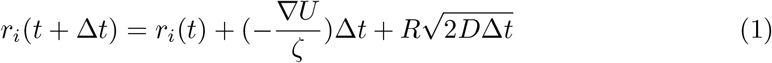

Where *r*_*i*_ is the position of the bead at time *t* in the direction *i* = x, y, or z, *R* is a random number drawn from a normal distribution with a mean of 0 and a standard deviation of 1, *ζ* is the drag coefficient, and *D* = *k*_B_*T/ζ* is the diffusion coefficient. The forces each bead experiences due to interactions with the surrounding polymer or target are captured in *U* where ∇*U* is a potential energy that combines contributions from connectivity, excluded volume, and binding. These are added together as *U* = *U*_sp_ + *U*_LJ_ + *U*_Bind_.

Connectivity along the polymer chain is controlled by harmonic springs with the equation:

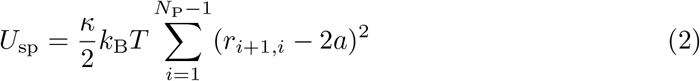

where *r*_*ij*_ is the distance between polymer beads, *N*_P_ is the degree of polymerization of the polymer in beads, *a* is the radius of a simulation bead, and *κ* was chosen to be 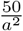, a value sufficiently large enough to prevent the polymer from stretching apart under normal Brownian forces.

A generic Lennard-Jones potential was applied to control excluded volume and implicit solvation according to:

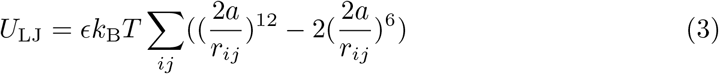

Where *i* and *j* represent two different bead indices and the value of *ϵ* can be adjusted to control the solvent quality and non-specific interactions between beads. We also use this Lennard-Jones potential to create non-specific attraction between target binding proteins and polymer beads as shown in Fig 2A. This generic potential could represent attraction due to hydrophobicity or Van der Waals. The range of the non-bonded potential has been shown to change the liquid-solid phase boundary in colloidal systems, but the gas-liquid boundary was not qualitatively changed [21]. Therefore, because this work focuses on the gas-liquid boundary in relatively dilute systems, we could add a screened electrostatic potential (a shorter range interaction) but do not expect this to qualitatively change our results. Unless otherwise specified, we chose *ϵ*_PP_ = 0.41 to mimic polymer configurations in a theta solvent [22] as shown in Fig 3A. For protein targets binding with valence-limited lock and key bonds, we used polymer target potential *ϵ*_PT_ = 0.1 to mimic a good solvent and separate non-specific and specific binding interactions. Between the targets themselves, *ϵ*_TT_ was varied from 0.5 to 3.0 to capture a range of target protein solubilities. The specific values of *ϵ*_TT_ are specified in later requisite sections.

**Fig 1.**
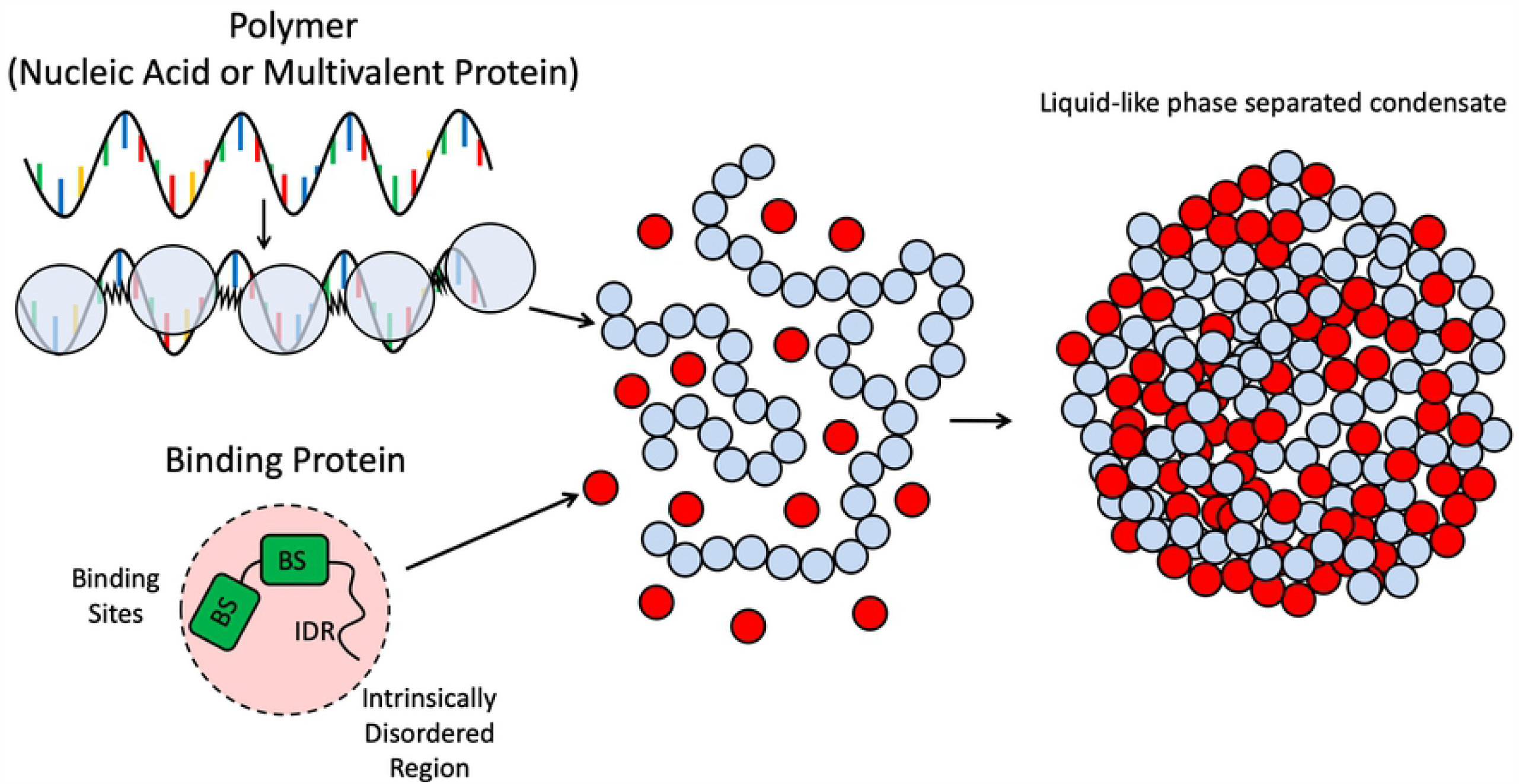
Depiction of simulation scheme. Polymers are represented by spherical beads (light blue) connected by harmonic springs. These polymers could represent either nucleic acids or long modular binding proteins found in biocondensates. Each polymer bead has a single binding ligand. Target binding proteins are represented as spherical beads (red) and can have multiple binding sites (BS) depicted as green blocks. These protein beads also encompass a intrinsically disordered region (IDR) that modulates their non-specific attraction to the polymer and between the proteins themselves. When the polymers and binding proteins are mixed together, they can undergo a phase transition into a condensed droplet.

**Fig 2.**
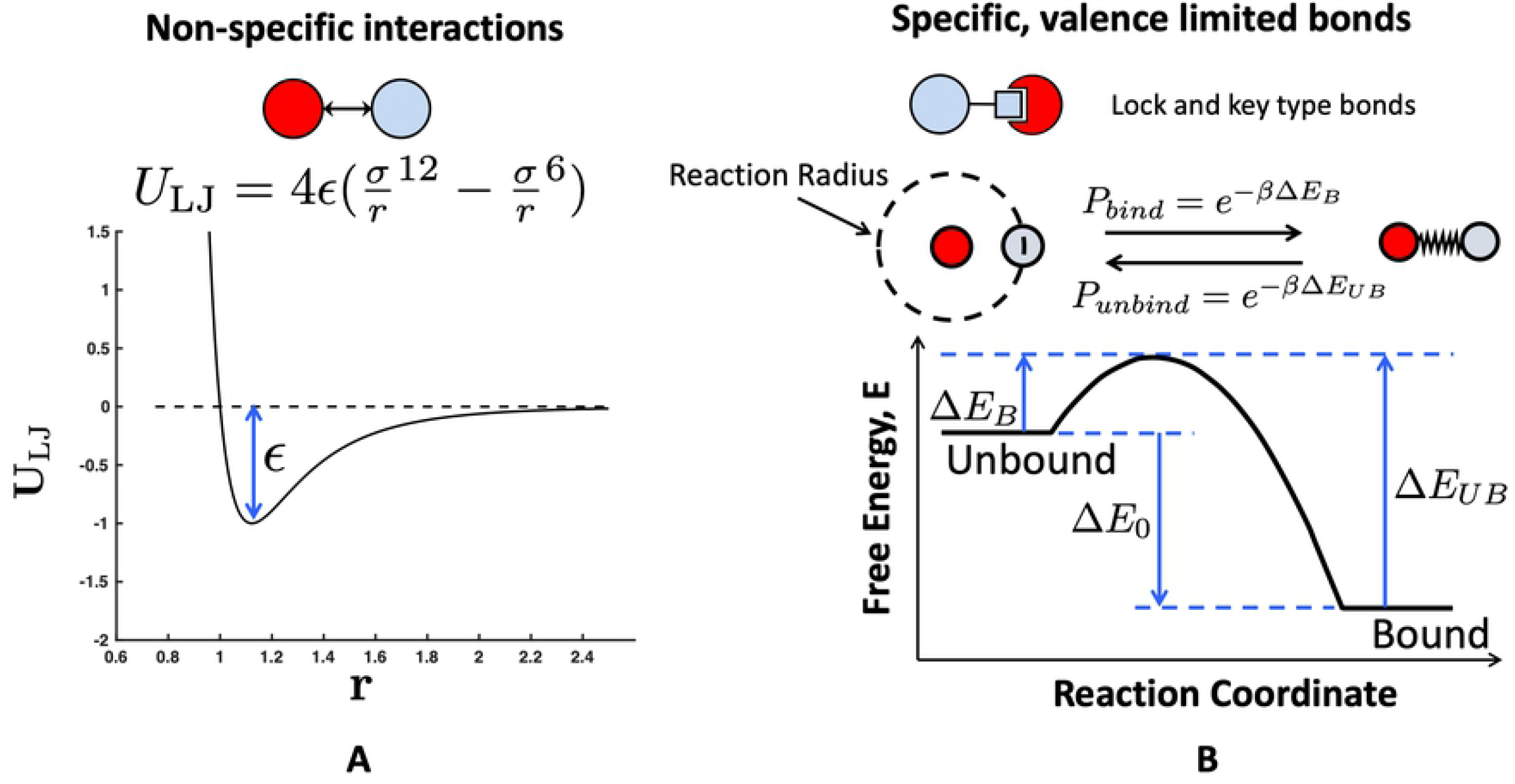
Two types types of protein-polymer interactions explored in this work. (A) Non-specific excluded volume interactions controlled by a Lennard-Jones potential. These potentials are not valence limited and are felt by any target or polymer bead in accordance with their distance apart *r*. (B) Specific, valence-limited, lock-and-key type binding. Polymer ligands and target protein binding sites interact when they are within a reaction radius that is dependent on the timestep. Within this reaction radius, they have a probability of binding *P*_B_ that depends on the depicted free-energy landscape. Once bound, the target and polymer bead are connected by a harmonic spring, and with some probability, *P*_UB_, can return to being unbound and interacting solely through a Lennard-Jones potential. This figure is adapted from Zumbro *et al*. with permission from Elsevier [19].

**Fig 3.**
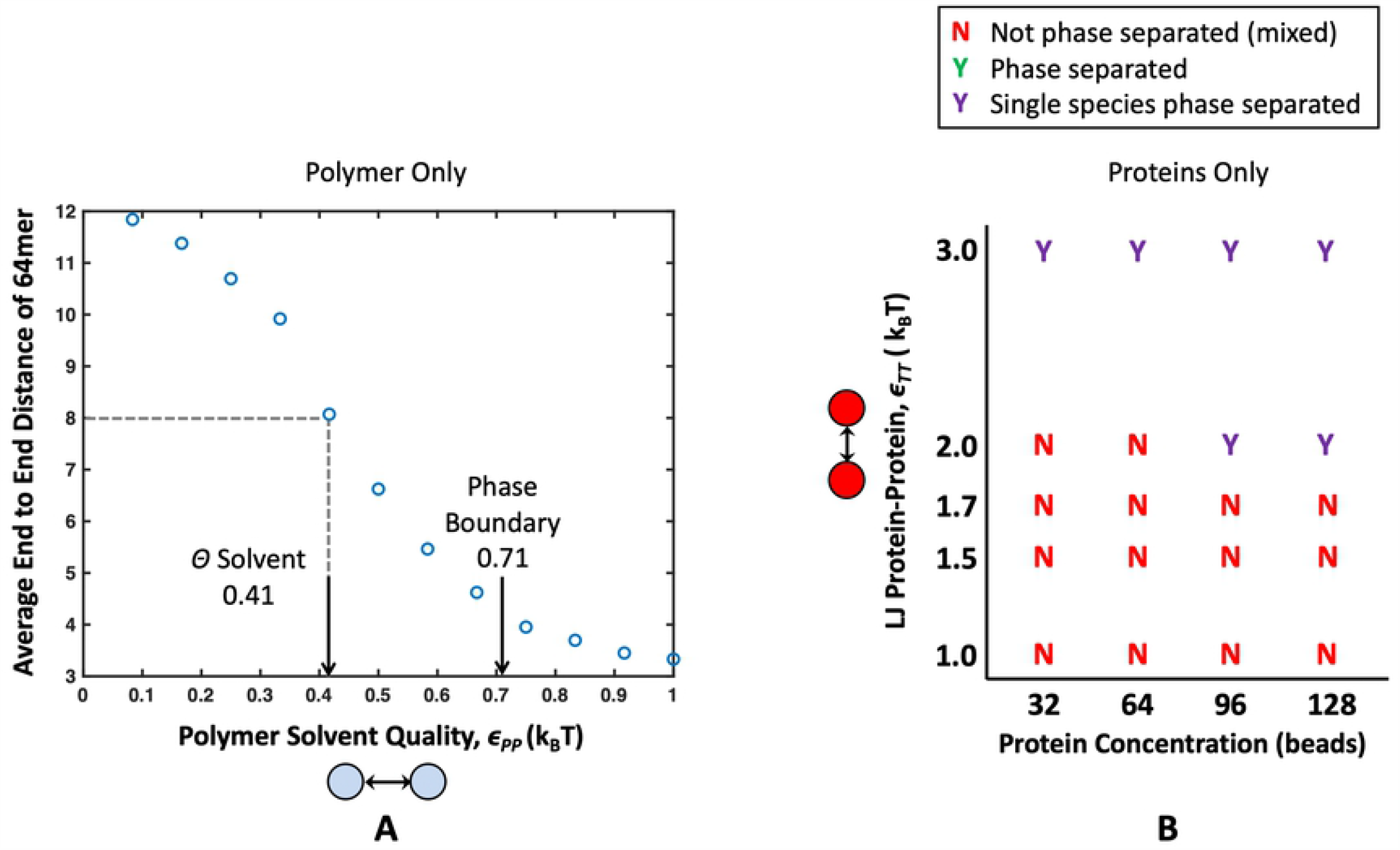
Properties of a single species alone, before mixing them together. Average end-to-end distance of a 64mer polymer under various Lennard-Jones attractions *ϵ*_PP_. The polymer behaves as it would in *θ* conditions, as a perfect random walk, when *ϵ*_PP_ = 0.41. *ϵ*_PP_ = 0.71 is highlighted with an arrow to denote the attraction at which four 16mer polymers aggregate into a single condensate. From this, we can see there is a region of poor solvent where polymers are collapsed but still soluble. (B) Phase diagram showing solubilities of binding proteins alone. When targets form a condensed phase without polymer, it is denoted with a purple “Y”, and when they do not form a condensed phase, it is denoted with a red “N”. From this chart, we see that all target concentrations tested are phase separated when *ϵ*_TT_ = 3.0, no target concentrations nucleate a condensed phase at *ϵ*_TT_ = 1.7, and only high target concentrations 96 and 128 targets phase separate at *ϵ*_TT_ = 2.0. This phase diagram will serve as a control for the effects of mixing polymers and target proteins. This figure is adapted from Zumbro *et al*. with permission from Elsevier [16].

The last type of interaction included is a reactive lock and key bond shown in Fig 2B, which represents our specific, valence-limited binding interaction between polymers and targets. To simulate this reactive binding, harmonic springs were turned on and off between the polymer beads and the targets to dynamically represent bonded and unbonded states. This was implemented using the prefactor Ω(*i, j*) multiplied by a harmonic potential as follows:

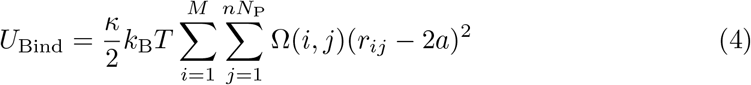

Where *M* is the valency of the target protein and *n* is the number of polymers in the simulation. Ω(*i, j*) = 1 when the *i*th binding site on the target is bound to the *j*th bead of the polymer, and Ω(*i, j*) = 0 when the target binding site or polymer bead is unbound. To control the probability of binding and unbinding, we use a piecewise function based on the energy barriers for the binding reaction from Sing and Alexander-Katz [23].

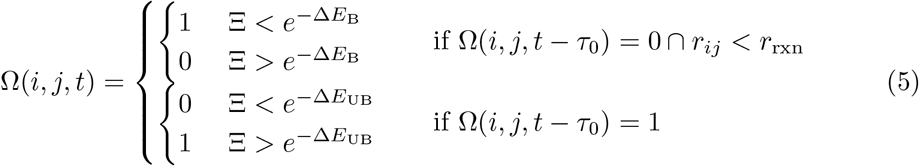

Here, Ξ is a random number between 0 and 1, Δ*E*_B_ is the energy barrier to bind normalized by *k*_B_*T*, and Δ*E*_UB_ is the energy barrier to unbind normalized by *k*_B_*T* as depicted in Fig 2B [23]. Without loss of generality, these energies are considered to be always positive, and the kinetics of binding are held constant by keeping Δ*E*_B_ at 0.5*k*_B_*T* so that binding is an accessibly frequent event. Increasing or decreasing the energy barrier will respectively slow or accelerate the kinetics of binding and unbinding equally, but not change the system’s thermodynamics. The thermodynamic drive of binding is controlled by varying Δ*E*_0_ = Δ*E*_B_ − Δ*E*_UB_. Binding becomes more favorable as Δ*E*_0_ is made more and more negative. This method is based directly on Sing *et al*. as well as others [23–26]. Binding reactions are evaluated every time interval *τ*_0_ = 100Δ*t*, where Δ*t* is the length of one timestep and *t* is the current time. The reaction radius *r*_rxn_ = 1.1 is the distance apart two beads would be if their surfaces were touching plus 0.1. Choosing 0.1 < (6*Dτ*_0_)^1*/*2^ allows time for a target that unbinds to diffuse out of the polymer radius of influence in *τ*_0_ and makes binding events independent [23]. We have applied the constraint that at any time, an inhibitor bead can only bind to one target binding site Σ_*j*_ Ω(*i, j, t*) ≤ 1, and a target site can only be bound to one inhibitor bead Σ_*i*_ Ω(*i, j, t*) ≤ 1. Competing reactions are sampled randomly. Note that we do not include the effect of forces in the breaking of the bonds; this is due to the fact that for forces on the order of *k*_B_*T/a*, this effect is negligible if the characteristic bond length is less than 1 nm. For reference, a discussion of the subject is given by Sing and Alexander-Katz [27]. The binding sites are isotropic to avoid unnecessary assumptions about binding site orientation and geometry in this coarse-grain model and to make the model more generally applicable. This means that binding is attempted and can be successful whenever the centers of an unoccupied polymer bead and unoccupied target bead are within a distance of *r*_rxn_ = 1.1.

This reactive binding scheme is applied with a free energy of binding per site of Δ*E*_0_ = −2, −4, and −6*k*_B_*T*. Each polymer bead contained a single binding site, and each target bead was given *M* = 1, 2, or 3 binding sites in order to capture the effects of changing binding valency. The simulation box had periodic boundaries with side length *l*_box_ = 41*a*. Because the box size is dependent on the size of the bead, we can convert the target concentration from beads per box to Molar by assuming a binding protein radius. Assuming the target bead radius to be *a* = 2.5 nm results in a target concentration of approximately 1.6 *µ*M per bead (e.g., 64 targets per box is approximately 102 *µ*M, 96 targets ≈154 *µ*M). Similarly, by assuming the size of a target bead radius to be approximately *a* = 2.5 nm, and using Langmuir adsorption theory, we can convert this Δ*E*_0_ = −2, −4, and −6*k*_B_*T* binding energy into a dissociation constant in Molar, resulting in a monovalent binding affinity of approximately *K*_D_ = 0.8, 0.1, and 0.02 mM, respectively. Details of this conversion are shown in the supporting information. This monovalent binding affinity is well within the weakly binding mM to *µ*M affinity range of some monovalent protein-protein and RNA-protein binding found in biocondensates [5, 28, 29].

The potentials are applied over the timestep 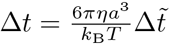 where 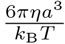 is the characteristic monomer diffusion time or the time that it takes a bead to diffuse its radius *a* and the dimensionless timestep is 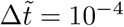. These equations are all made dimensionless by scaling energies by thermal energy *k*_B_*T*, lengths by bead radius *a*, and times by the characteristic diffusion time 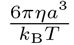. Results are averaged over the last quarter of the total simulation time and over at least 10 different runs, with a typical system energy profile over time shown in supporting information S2 Fig.

The simulation code is available by request.

## Results and Discussion

In biocondensates, components can often phase separate by themselves, but interactions with another species can cause phase separation at different energies or concentrations [3, 14, 15, 30]. Previous research has also shown that altering stoichiometry and the mismatch in concentrations between receptors and ligands can lead to switchlike control of biocondensate formation [8]. Therefore, we wanted to explore the how the phase boundary of our system changes with the ratio of polymer to target, and subsequently how this dependence is modulated by the multivalent polymer properties. As a control, we first ran simulations of a single species, only targets and only polymers, to understand their solubility before mixing at different inter-polymer and inter-target attractions. Results for simulations with polymers only and targets only are plotted in Figs 3A and B, respectively. Polymers with a degree of polymerization of 64 beads, behaved as freely-jointed random walks, characteristic of *θ* solvent conditions at *ϵ*_PP_ = 0.41, and consistent with previous literature [22]. Four 16mer polymers were seen to condense into a single droplet at *ϵ*_PP_ *>* 0.71. This means that, at the polymer concentrations used throughout our simulations, there is a region of *ϵ*_PP_ where polymers can experience poor solvent quality but not phase separate on their own. Throughout this work, we will consider polymers with *ϵ*_PP_ ≤ 0.58, meaning that in all results discussed below, the polymers do not phase separate on their own.

We also consider the potentials necessary to phase separate the targets alone at our simulation concentrations (32, 64, 96, and 128 targets). By themselves, target concentrations of 96 and 128 formed a condensed phase at *ϵ*_TT=2.0_, all target concentrations nucleated a condensed droplet at *ϵ*_TT_ = 3.0, and no targets phase separated at *ϵ*_TT_ = 1.7. These inter-target energies bounded the parameter space for our simulations where polymers and target proteins were mixed together. In later phase diagrams, energy and concentration regions where targets can phase separate on their own are shaded with a purple background.

### Valency and Affinity of Specific Lock and Key Bonding

To compare our system to previous computational studies, we first investigated the effects of valency and binding site affinity on the LLPS of multivalent polymers and targets with specific, valence limited binding interactions. To do so, we placed four polymers with a degree of polymerization *N*_P_ = 16 beads in *θ* solvent with 32, 64, 96, and 128 binding protein targets. Monovalent, divalent, and trivalent targets were simulated with the 3 different binding site affinities (Δ*E*_0_ = −2, −4, and −6*k*_B_*T*), and the resulting phase diagrams are shown in Fig 4. In order to determine if a system nucleated a stable condensed phase, used a combination of visual inspection, the Binder cumulant, and a collapse in the *R*_g_ as described in previous work [16]. We used visual inspection to look for the proportion of 10 runs in which a condensed droplet of targets and polymers persisted for the last quarter of the simulation time. Simulations in which a stable droplet formed more than 70% of the time are marked with a green “Y”, systems that formed a droplet in 60% of runs are marked with a yellow “Y”, and systems where less than 50% of runs formed a droplet are marked with a red “N” for no phase separation. Visually inspecting out simulations for dense droplets is very similar to the method used in Choi et al. where the authors determined phase separation by looking for the onset of density inhomogeneity [31]. We quantitatively confirmed phase separation by calculating the average energy of the last quarter of run time and using it to compute the Binder cumulant 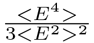. This quantity compares the ratio of the energy variance, which is equivalent to the specific heat of the system, to the average energy, and reaches a maximum at the phase transition [32]. Fig 5 shows an example of average energy and cumulant plots for a divalent protein at various concentrations with lock and key binding affinity of Δ*E*_0_ = −6*k*_B_*T*. By comparing the Binder cumulant along lines of constant target concentration, we confirmed the majority of our initial phase diagrams created through visual inspection. Areas where the cumulant predicts phase separation are shaded with a blue background in Fig 4 and later phase diagrams.

**Fig 4.**
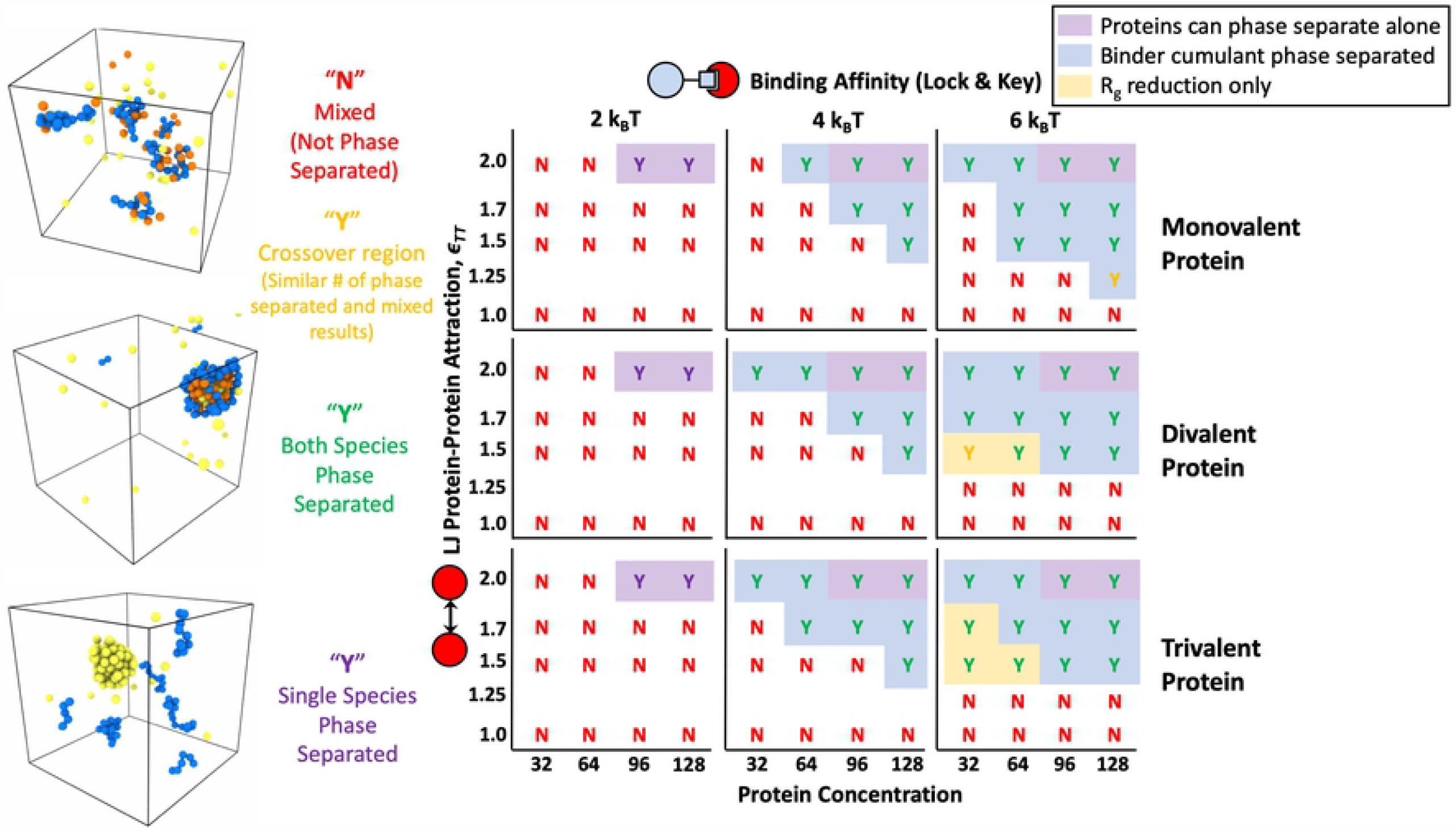
Phase diagram resulting from specific lock-and-key binding to four 16mer polymers. Results are shown for mono, di, and trivalent binding proteins with Δ*E*_0_ = 2, 4, and 6*k*_B_*T*. Letters and letter coloring were determined by visual inspection, with example renderings shown on the left of “Mixed” states labeled as a red “N” for no phase separation, fully phase separated systems with both polymers and proteins found in the condensed phase labeled with a green “Y” for yes phase separated, and purple “Y”s denoting systems in which a single species phase separated without the other such as the proteins condensing on their own. Yellow “Y”s denote systems in a the crossover region between phase separated and mixed where 60% of simulations showed a stable condensed droplet. Purple background shading denotes regions where pure protein simulations phase separated on their own without the help of the polymer. Blue background shading denotes the regions where phase separation was also indicated by Binder cumulant of the system energy. Yellow background shading denotes that aggregation of polymers into a droplet was indicated by a significant drop in the total *R*_g_ of the polymer system accompanied by a reduction in the *R*_g_ of individual polymers. Phase separation occurs at lower protein target concentrations and lower *ϵ*_*T T*_ as valency and binding affinity are increased.

**Fig 5.**
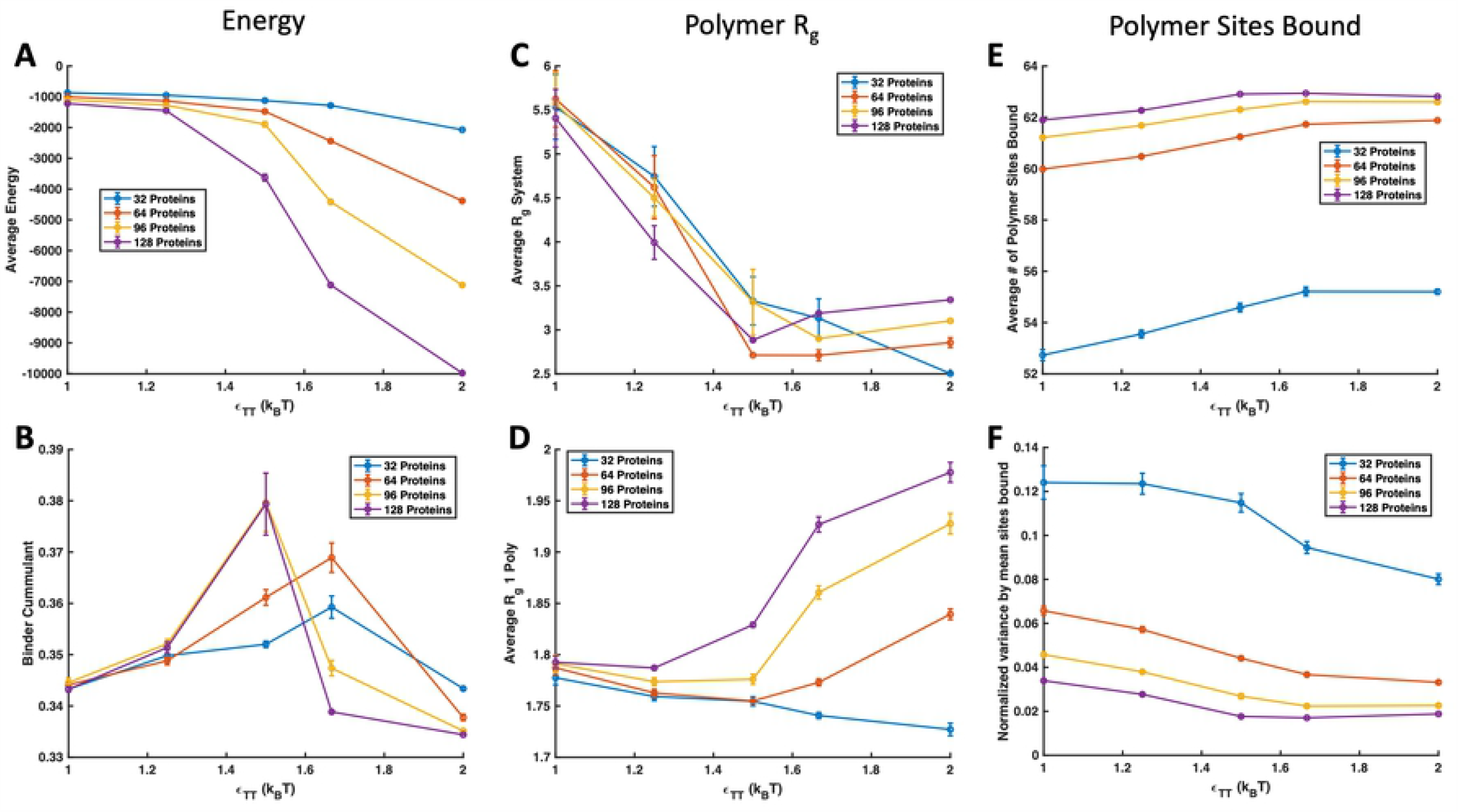
Example simulation properties. Examples of simulation properties for divalent protein targets with four 16mer polymers in theta solvent and Δ*E*_0_ = 6*k*_B_*T*, shown with lines of protein concentration. (A) Total average energy of simulation (B) Binder cumulant comparing average energy fluctuations to average system energy. A maximum in the Binder cumulant corresponds to a phase boundary. (C) *R*_g_ of all polymers in the system. A large reduction in system *R*_g_ signifies that all four polymers aggregated into a single body. (D) Average *R*_g_ of individual polymers across the simulation time. A reduction in *R*_g_ signifies a change in effective solvent conditions for the polymer as a result of complexation with binding proteins. After a critical concentration of protein binding is achieved, the polymers swell if the simulation isn’t protein concentration-limited. (E) Number of polymer sites bound with a maximum of 64. A plateau in sites bound occurs when a protein-polymer droplet is formed because the local concentration of protein targets reaches a maximum. (F) Variance in polymer binding sites occupied normalized by the average number of sites bound. The variance also plateaus when a condensed droplet is formed due to the smaller fluctuations in local concentration of proteins near the polymer in a liquid droplet.

We found that the Binder cumulant did not fully predict conditions in which we saw condensed droplets during visual inspection, so we also calculated the radius of gyration *R*_g_ of the polymers individually and all together to capture when the polymers themselves showed aggregation and collapse. Methods of measuring aggregation through *R*_g_ have been used in previous computational work on phase separation of biocondensates [6]. When four 16mer (*N*_p_ = 16) polymers come together in *θ* conditions to form a liquid droplet, without considering any swelling from binding proteins, they should have a similar *R*_g_ to a polymer with *N*_p_ = 64. In the ideal case, the size of a random coil polymer with *N*_p_ = 64 is 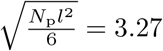 where *l* is the diameter of a bead 2*a* [33, 34]. In the example shown in Fig 5C, all target concentrations show clear polymer aggregation with *R*_g_ ≤ 3.27 at *ϵ*_TT_ = 1.5 and show an reduction in the individual polymer sizes around *ϵ*_TT_ = 1.25 in Fig 5D. This aligned well with our observation of condensed droplets at energies lower than shown by the Binder cumulant. Values that showed system-wide polymer aggregation and individual polymer *R*_g_ reduction but not a phase transition using the variance in system energy are shaded with yellow in Fig 4 and subsequent phase diagrams. For additional insight, we have also included plots of the average and variance in the number of polymer sites occupied by binding proteins in Fig 5E and F, respectively. There is an increase in average sites bound and a decrease in the variance of polymer sites occupied with increasing *ϵ*_TT_. The number of sites bound and the variance start to plateau upon polymer/target condensation because the local concentration of proteins within reach of the polymer reaches a maximum in a the condensed phase.

We expect that the discrepancy of the polymer-target binding system showing aggregation and individual polymer *R*_g_ reduction at lower energies than the Binder cumulant predicts a phase transition is the manifestation of two different transitions. As we increase the target-target attraction, bound targets create an effective interaction between polymers so that the polymers behave as if they are in poor solvent. This leads to a first transition where the polymers aggregate and a condensed polymer droplet or small polymer gel forms. This is similar to results reported by Harmon et al. where a wider set of species concentrations resulted in gels than phase separated condensates [6]. This polymer aggregation does not manifest in the Binder cumulant because intra-target interactions are stronger than intra-polymer interactions and dominate the mean energy and specific heat of the system. Although the polymers have aggregated during the first transition, the targets are still relatively disperse, overwhelming the energy variance and the Binder cumulant arising from polymer aggregation. As the intra-target attraction *ϵ*_TT_ is further increased, we see a second transition captured in the energy and cumulant because the polymer clusters nucleate a condensed target phase. If we imagine the polymer as RNA, this result is consistent with the idea that condensed RNAs can act as scaffolds for nucleating condensates [15].

By returning to explore our phase diagram in Fig 4, we now see that the addition of the binding polymer leads to lower target solubilities for all target valencies studied for both Δ*E*_0_ = −4*k*_B_*T* and −6*k*_B_*T*. The targets phase separate more at more easily when binding to a multivalent polymer. The boundary for target phase separation in the absence of polymer is highlighted with a purple background. As the affinity of the specific binding sites increase, the phase boundary shifts down to weaker *ϵ*_TT_ and lower target concentration. Although less dramatic, a similar shift is seen with target valency. As target valency increases, phase separation occurs at lower target attraction *ϵ*_TT_ and target concentration. This result matches well with previous simulations and experiments [5, 6, 35, 36] and demonstrates that condensed droplets and similar behavior can be seen in much smaller systems than previously reported. This suggests that condensates can from droplets much smaller than can be seen through a microscope, and large condensates might grow through coalescence of these smaller droplets.

We also explored phase diagrams through the lens of polymer valency or length. Fig 6 shows phase diagrams for divalent targets binding through reactive specific binding with multivalent polymers. In these simulations, the number of polymer beads was kept at a constant concentration, but the connectivity of the polymers was changed to measure the affect of polymer valency and degree of polymerization. Simulations contained sixteen 4mer polymers, four 16mer polymers, or one 64mer polymer. Consistent with previous results on increasing valency leading to droplet formation at lower concentrations and binding affinities, we also saw phase separation at lower energies for polymers with higher degrees of polymerization [5, 6, 35, 36]. The lowering of the phase separation boundary is more drastic for Δ*E*_0_ = −6*k*_B_*T* when the polymer length is increased from *N*_p_ = 4 to *N*_p_ = 16 than when the polymer length is increased from *N*_p_ = 16 to *N*_p_ = 64 even though both scenarios reflect a 4X increase in length. As explored in our previous work, binding affinity of a linear multivalent polymer to a smaller protein target is significantly affected by the entropic cost of forming polymer loops when binding to the target divalently [19]. As degree of polymerization increases, longer loops are able to form, adding combinatorial entropy and increasing binding affinity. However, at some critical loop size, the maximum possible loop length is limited by the configurational entropy loss of forming the loop and not limited by the polymer length. Polymers longer than the critical loop length see limited increases in avidity with longer degrees of polymerization. In these phase separation simulations, this results in a relatively small increase in binding avidity when *N*_p_ is increased from 16 to 64 (and a correspondingly smaller change in the phase boundary) than when *N*_p_ is increased from 4 to 16.

**Fig 6.**
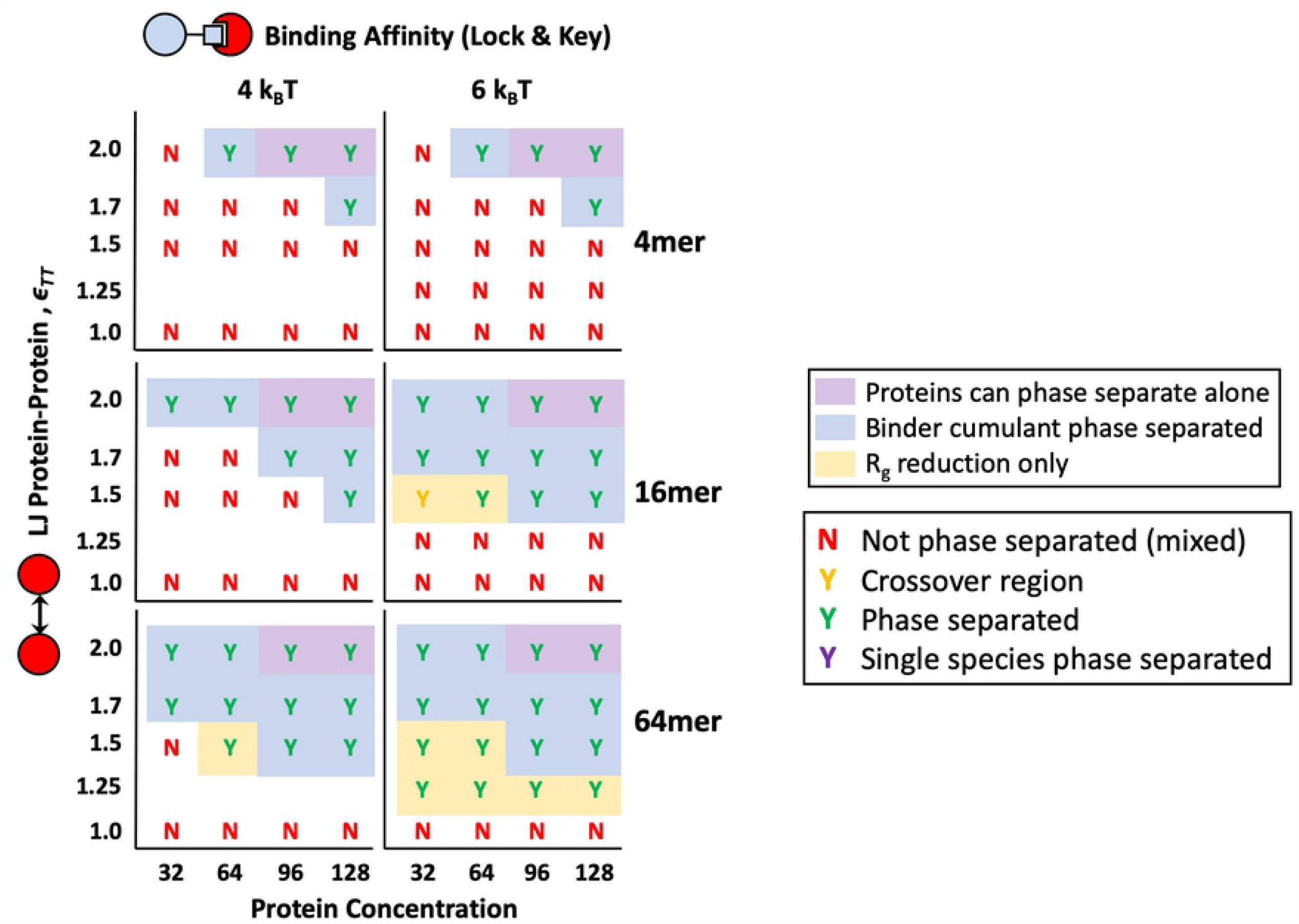
Phase diagrams for polymers of different of degrees of polymerization. *N*_p_ = 4 (Top Row), 16 (Middle Row), and 64 (Bottom Row) with reactive, specific binding affinities Δ*E*_0_ = −4 and −6*k*_B_*T*. All simulations are for divalent proteins targets and theta solvent for the polymer. Letter color coding and area shading have the same meanings as described in Fig 4.

### Solvent Quality

Previous research showed that native intrinsically disordered protein (IDP) linkers on multivalent polymers can be swollen, theta condition freely random walk, or collapsed chains [6]. This same study showed that highly solvated or swollen polymers initiated gelation without phase separation and theta polymers led to phase separation with gelation [6]. Although not explored in prevously, LLPS in poor solvent quality is interesting because 30% of IDPs in the aforementioned study were found to have negative solvation volume. The effects of multivalent polyemrs in poor solvent are not immediately obvious because a condensed polymer’s binding sites may be less available for target binding, or the polymer could phase separate on its own without the targets. Also, unlike good and theta solvent, polymers in poor solvent effectively have multivalent binding interactions with both themselves and their binding protein targets. Using a Lennard-Jones potential and Brownian Dynamics, we show that there is a window of poor solvent conditions where polymers are soluble on their own, but can nucleate condensed droplets in the presence of binding targets.

Here, we again placed four 16mer polymers in a box with divalent targets with increasing target concentration and *ϵ*_TT_. If four 16mer polymers are simulated in a box by themselves, they precipitate out of solution at approximately *ϵ*_PP_ = 9*/*12, so we tested energies between *ϵ*_PP_ = 5*/*12 and 8*/*12 where the 16mer polymers were collapsed but still soluble. Resultant phase diagrams for theta and poor solvents with *ϵ*_PP_ = 6*/*12 and 7*/*12 are shown in Fig 7.

**Fig 7.**
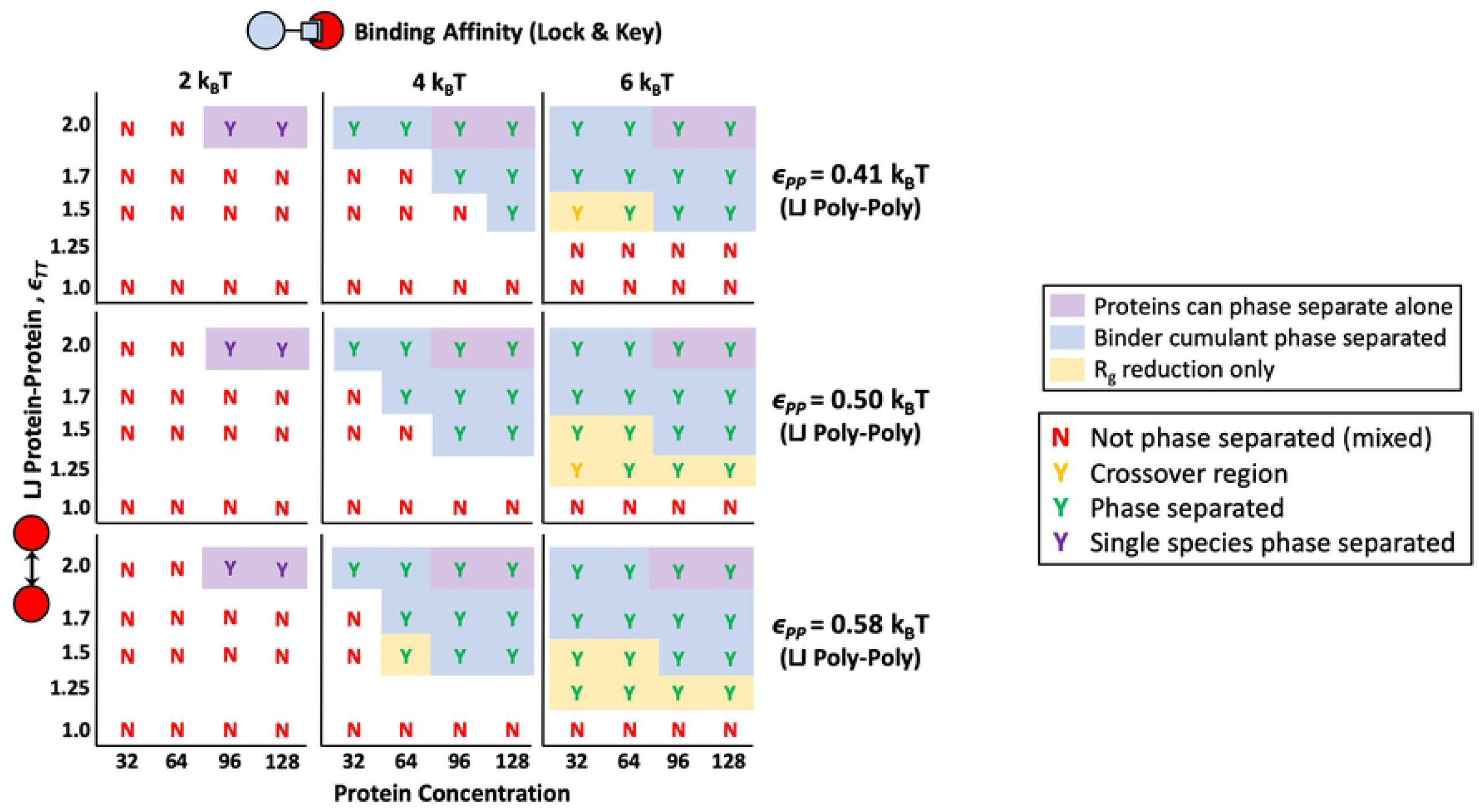
Phase diagrams comparing theta and poor solvent. Phase diagram of simulations comparing the behavior of four 16mer polymers with lock and key binding to divalent protein targets in theta solvent (*ϵ*_PP_ = 5*/*12) to two types of poor solvent (*ϵ*_PP_ = 6*/*12 and 7*/*12). Polymers do not phase separate on their own at any values of *ϵ*_PP_ tested. Lettering color codes and shading follow the same key as Fig 4, with “Y”s indicating “yes” phase separation occurred and “N”s representing “no” phase separation occurred. In poor solvent, phase separation occurs when polymers are mixed with binding targets at lower *ϵ*_TT_s and protein concentrations than theta solvent.

In the case of poor solvent, even though collapsed polymers have less available volume for targets to bind in, polymers phase separate at lower target-target attraction and target concentrations than polymers in theta solvent. Results from our general collapsed polymer model demonstrate that a slight decrease in solvation of the polymer may trigger phase separation when the polymer is mixed with a corresponding binding target. This phenomenon can still happen when the decrease in solvent quality is not enough for the polymer to precipitate on its own. For example, changes in polymer sequence, post-translational modifications, or binding to a protein or small molecule that lowers the effective solvent quality for a multivalent polymer could all cause phase separation at lower binding energies, concentrations, and target-target attractions. Our results demonstrate how decreasing solvent quality lowers the phase boundary for systems with multiple components, building on recent work from Martin et al. that showed pure multivalent polymers phase separate at higher temperatures when they have more attractive self-interactions [37].

### Non-specific binding interactions

In addition to multivalency through valence-limited specific binding sites, we wanted to consider any differences in phase separation behavior associated with non-specific interactions such as charge or hydrophobicity. Non-specific interactions are commonly believed to add additional valency to lock-and-key binding through the promiscuous interactions of IDRs on binding proteins such as FUS, TDP43, and hnRNPA1 [13, 38]. To isolate the effects of non-specific interactions on nucleating a condensed phase, we turned off our reactive binding scheme and exclusively applied a more attractive Lennard-Jones attraction between the polymers and the targets *ϵ*_TP_ as shown in Fig 2A. We again placed four 16mer polymers in theta solvent with various target concentrations and *ϵ*_TP_s. The phase behavior for *ϵ*_TP_ = 0.75, 1.0, 1.25, and 1.5*k*_B_*T* is shown in Fig 8. Results for a single longer polymer with degree of polymerization *N*_P_ = 64 are also included for comparison.

**Fig 8.**
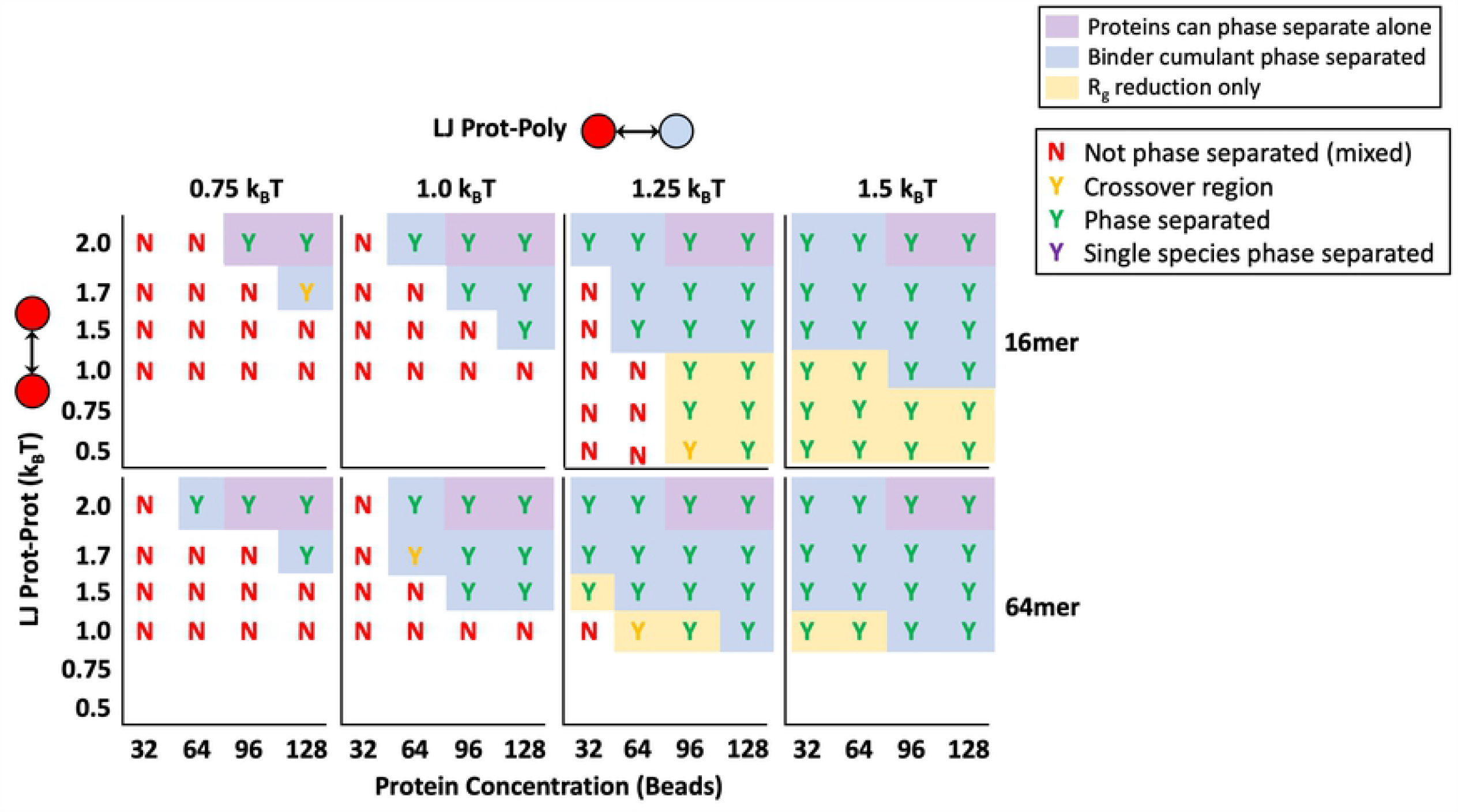
Non-specific binding phase diagram. Phase diagram for proteins binding to polymers through a non-specific Lennard-Jones potential. Diagrams use the same key described in detail in Fig 4 where phase separated systems are marked with a “Y” for “yes” and not phase separated systems are marked with an “N” for “no”. Results are shown for simulations with increasing polymer binding attractions *ϵ*_TP_ from left to right (columns) with four 16mer polymers (Top) and one 64mer polymer (Bottom).

With a generic Lennard-Jones attraction between target proteins and the polymer, phase separation occurs at much lower potential energies than with specific, reactive binding. For example, all target concentrations and solubilities as low as *ϵ*_TT_ = 0.5*k*_B_*T* showed droplet formation at non-specific attraction *ϵ*_TP_ = 1.5*k*_B_*T*, but with the highest reactive binding energy Δ*E*_0_ = −6*k*_B_*T*, only targets with *ϵ*_TT_ ≥ 1.5*k*_B_*T* formed droplets. This huge shift in the phase boundary toward lower energies as we change from specific lock-and-key to non-specific bonding is likely because the non-specific interaction has a much higher valency that is only limited by the maximum number of neighbors. The limited valence of lock-and-key type bonds creates competition for sites between bound and unbound target neighbors, reducing the influence of the polymer on the targets. This reduction in binding due to competition for high affinity sites is discussed further in our previous work on multivalent binding site patterns [39]. The promiscuous nature and high effective valency of non-specific attraction reduces competition and allows polymers to interact with more targets simultaneously. This results in polymer-target phase separation occurring at much lower attractions and concentrations. Therefore, non-specific interaction energies are a very sensitive dial for controlling phase separation and polymer modifications that change the non-specific interactions such as hydrophobicity or charge between the polymer and targets will have a more significant impact on the phase boundary than alterations to specific binding sites.

Still, valency and affinity of specific bonds can also change the phase boundary, although relatively large changes in these characteristics correspond to small changes in the LLPS boundary. Consequently, with a combination of non-specific and lock and key binding, the LLPS of multivalent polymers and targets can be precisely controlled. An example of using both non-specific and specific binding is shown in Fig 9. A small non-specific attraction *ϵ*_TP_ = 5*/*12*k*_B_*T* was applied in addition to specific divalent binding with Δ*E*_0_ = −2, −4, and −6*k*_B_*T*. The addition of the non-specific attraction provides access to phase separation at lower intra-target attractions, previously inaccessible through purely lock-and-key binding or at such a low specific polymer attraction. The non-specific binding makes the phase separation energy barrier accessible, and the specific, valency limited bonds can be used to tune the exact boundary. This is helpful because we showed above that the resultant phase boundary from specific bonds is less sensitive to changes in binding affinity or valance than the non-specific attraction. We speculate that some reliance on the insensitivity of specific bonding could help biology to reduce the number of aberrant phase transitions caused by unintentional changes in binding sites.

**Fig 9.**
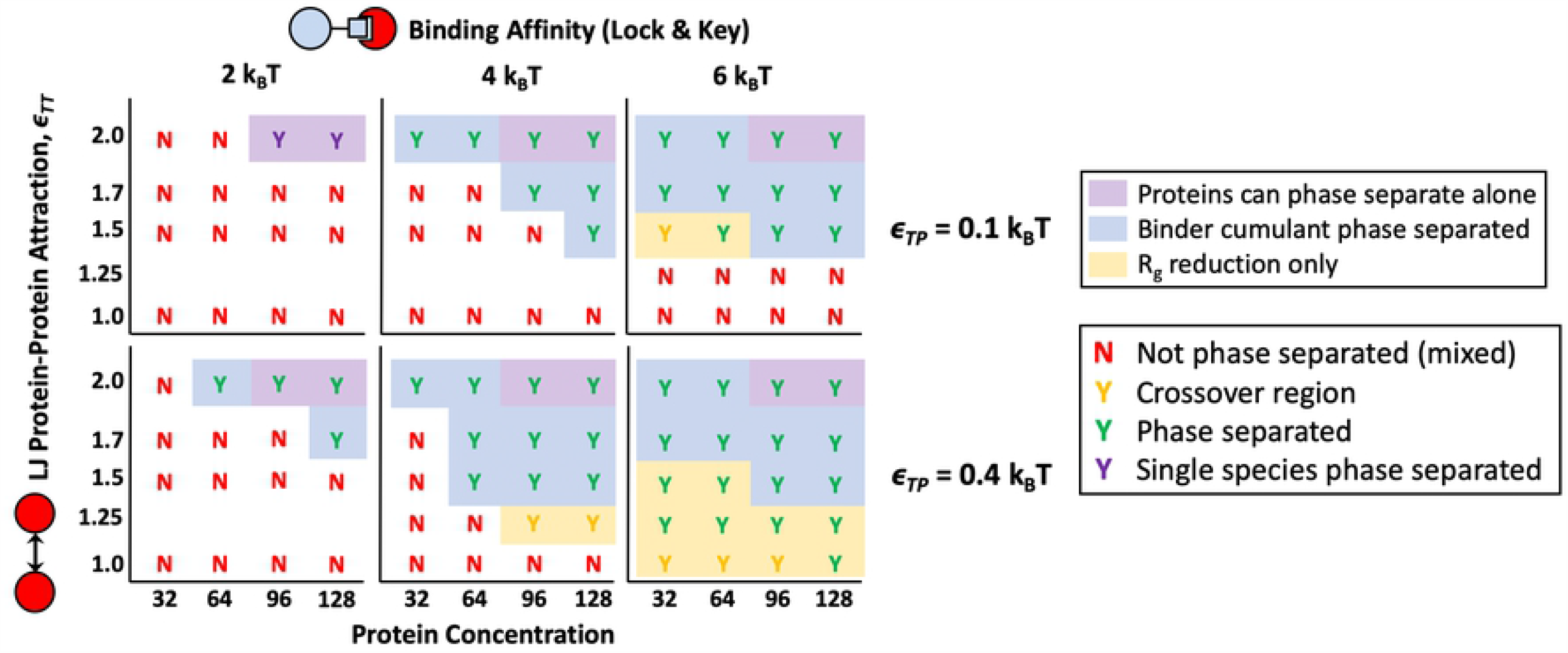
Phase diagrams with both non-specific and specific binding. Phase diagram of targets and polymer with both non-specific binding affinity *ϵ*_TP_ = 5*/*12 and specific lock and key binding (Bottom Row) compared with polymers that have lock and key binding but almost no non-specific attraction to the targets *ϵ*_TP_ = 5*/*12 (Top Row). Protein targets in these simulations are divalent. Diagrams use the same key described in detail in Fig 4 where phase separated systems are marked with a “Y” for “yes” and not phase separated systems are marked with an “N” for “no”. Results are shown for simulations with specific polymer binding attractions Δ*E*_0_ = −2, −4, and −6*k*_BT_ from left to right (columns) with four 16mer polymers.

### Condensed phase organization

Biocondensates often show microphase separation within the condensed phase [2, 3, 35, 40]. This disorder to order transition is a well known phenomenon in polymer physics with block copolymers where self-assembly can be controlled by the interaction (*χ*) between the two polymer block types [41]. We expected the same to be true for polymer-target assemblies in LLPS. When attraction between binding proteins themselves is higher than attraction to the polymer, condensates should undergo microphase separation where polymers surround a condensed target phase. When attraction between targets is similar to target-polymer attraction, the condensed phase will remain mixed. If the targets are highly attracted to the polymer, and not attracted to themselves, we might also see the case where the polymer is condensed in the center of the droplet with targets decorating the outside of the condensed phase.

First, we explore inducing droplet order by changing the self-interactions of a single species such as the binding proteins. Modulating self-interactions or solvation volume can be thought of as changing the surface tension of the liquid target phase. Changes in droplet organization due to surface tension or interactions of polymers with solvent were previously explored in a 4 component system with 3 types of equal size binding polymers and solvent. They found that swollen multivalent polymers could lower the surface tension of similar less-solvated polymers and induce a shell-core structure seen in some RNP bodies [40]. In Fig 10, we show that a similar ordering can be induced through surface tension in our asymmetric valency/size 3 component system, where the 3 components in our system are multivalent polymer, smaller binding proteins, and implicit solvent. If the target-polymer attraction is held constant, the condensates go through a demixing transition as the target-target attraction increases from *ϵ*_*T T*_ = 1.0*k*_B_*T* to 2.0*k*_B_*T* (moving in the vertical direction up the phase diagram in Fig 10). Similar changes in droplet organization controlled by the excluded volume of the binding proteins is also seen for the specific lock-and-key binding polymers in Fig 11.

**Fig 10.**
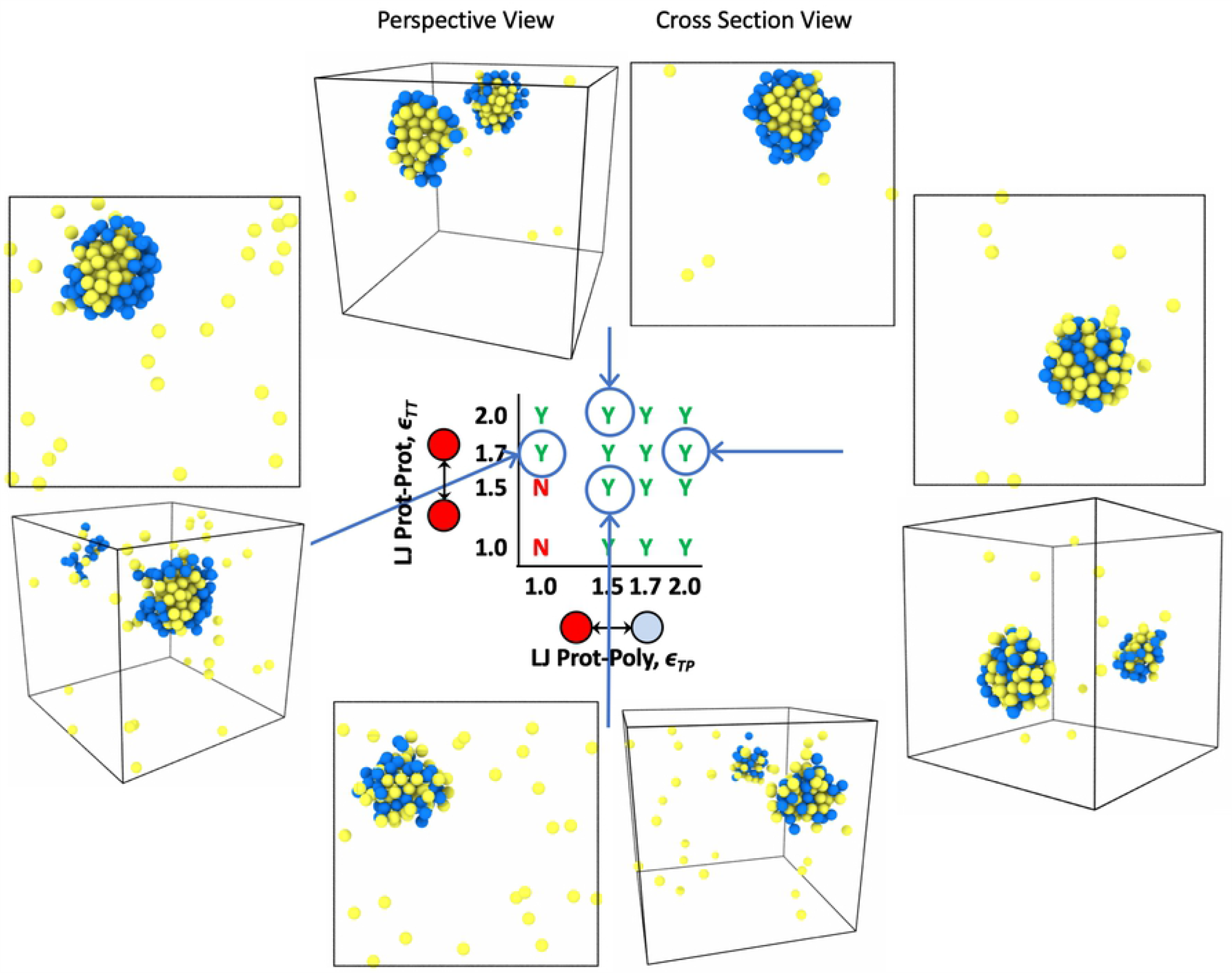
Renderings of droplet order with non-specific binding. Simulation renderings depicting ordered and mixed droplets with a cross section view through the middle of the droplet and a perspective view showing the inside and outside of the droplet. Polymer beads are blue and protein target beads are yellow. Results shown are for simulations with non-specific binding to four 16mer polymers in theta solvent and 96 target binding proteins. Note that the x-axis on this phase diagram is now protein-polymer affinity *ϵ*_TP_ in units of *k*_BT_ and the y-axis is still the intra-protein attraction *ϵ*_TT_ seen on previous phase diagrams also in *k*_B_*T*. By moving vertically down the phase diagram from *ϵ*_TT_ = 2.0 to 1.5 the droplet morphology goes from ordered to mixed due to changes in surface tension of the liquid protein phase. The droplet also goes from ordered to mixed as we move from left to right across the phase diagram from *ϵ*_TP_ = 1.0 to 2.0 due to increasingly favorable protein-polymer interfacial energy *χ*.

**Fig 11.**
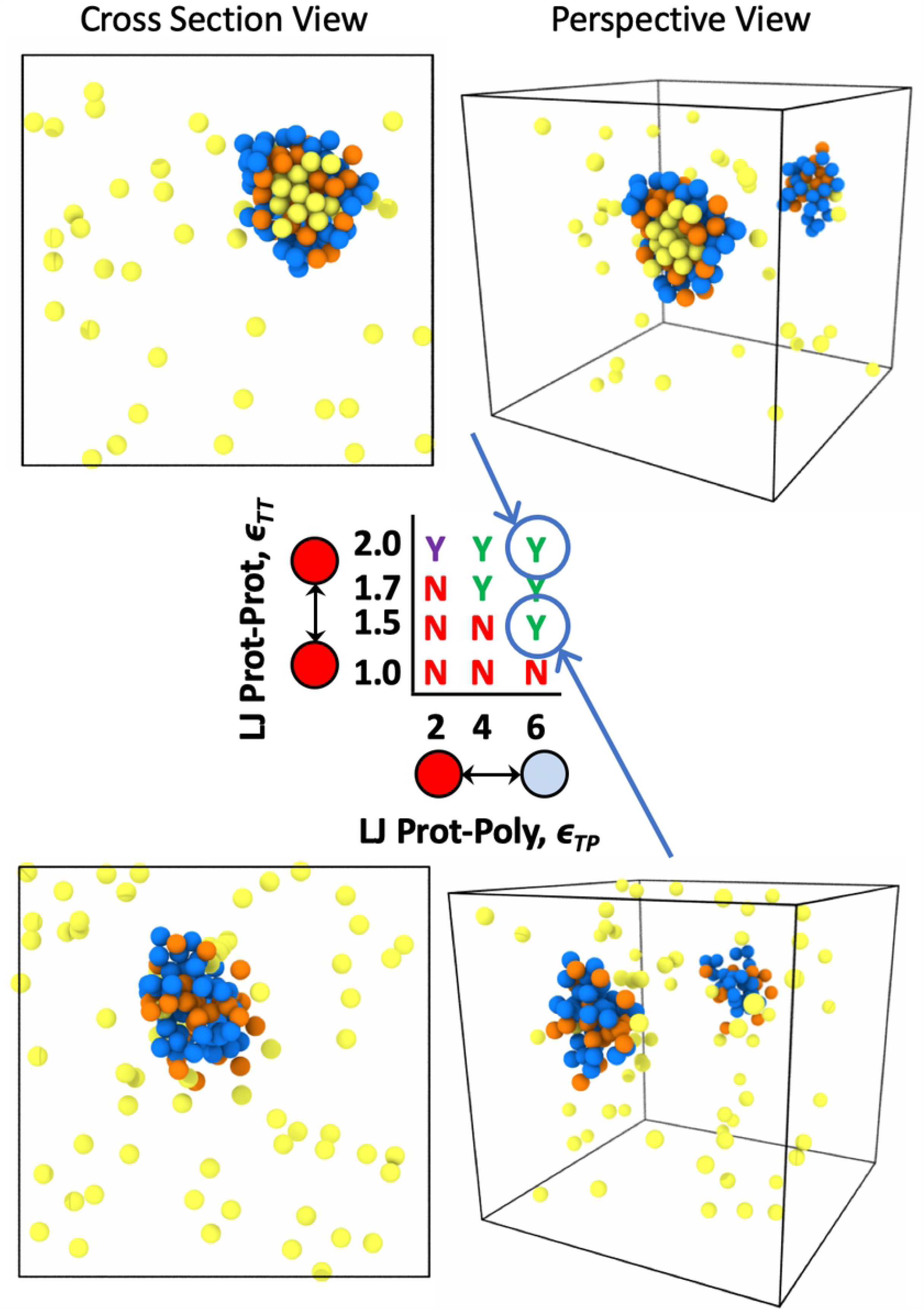
Renderings of droplet order with specific binding. Simulation renderings depicting ordered and mixed droplets with a cross section view through the middle of the droplet and a perspective view showing the inside and outside of the droplet. Polymer beads are blue, unbound protein targets are yellow, and bound protein targets are orange. Results shown are for simulations with specific binding to four 16mer polymers in theta solvent and 96 divalent target binding proteins. Note that the x-axis on this phase diagram is now protein-polymer affinity Δ*E*_0_ in units of *k*_B_*T* and the y-axis is still the intra-protein attraction *ϵ*_TT_ in *k*_B_*T*. By moving vertically down the phase diagram from *ϵ*_TT_ = 2.0 to 1.5 we also see the droplet morphology change from ordered to mixed due to changes in surface tension of the liquid protein phase.

In addition to ordering due to surface tension, which we control by modulating *ϵ*_*T T*_, we also see demixing caused by changes in non-specific binding affinity between the polymers and targets. Changing the non-specific binding affinity *ϵ*_*T P*_ is more akin to inducing order/disorder through modulating the *χ* parameter. Looking at droplet order in Fig 10, while moving across the phase diagram from left to right, it is clear in the simulation renderings that the system goes through and order-to-disorder transition as the target-polymer attraction increases from *ϵ*_*T P*_ = 1.0*k*_B_*T* to 2.0*k*_B_*T* for constant *ϵ*_*T T*_ = 1.7*k*_B_*T*. In this case, when the bonding energy or attraction between the targets and polymers are very strong, the targets prefer to associate with the polymer equal to or more than associating with themselves and subsequently decorate the polymer as much as possible. This results in the polymer becoming fully mixed with the binding targets. In contrast, when the target attraction to the polymer is less than the amount targets like to associate with themselves, the targets prefer to surround themselves with only target neighbors, and the polymer is pushed to the outside of the droplet. These results show that modifications made to targets that impact binding sites or non-specific attraction to a multivalent polymer and not just excluded solvent volume can change droplet ordering. For example, if the target is a protein and the polymer is RNA, a modification to the IDR region on the protein or a base pair on the RNA that could lead to changes in droplet ordering, if the altered sequence impacts the attraction between the protein and the RNA.

### Kinetics inside the droplet

Our simulation methods also allow us to study the diffusion and dynamics of condensed phase species which are important to understand biocondensate function and diseases associated with altered droplet dynamics [14, 17, 42, 43]. To examine the mobility of binding proteins and polymers in droplets, we looked at both the mean squared displacement (MSD) and the neighbor persistence. We define neighbor persistence as the average number of neighbors that remained the same over a time interval, where beads were considered neighbors when their centers were within 2.5*a* [44]. The rate at which the neighbor persistence goes to zero provides a measure of how quickly the targets are exchanging with the dilute supernatant phase. A high neighbor persistence or slow decay rate means that a target in the droplet maintains many of its neighbors over a long period of time, suggesting a more solid-like phase. A fast decay in neighbors to zero signifies that, in a short amount of time, the target became surrounded by an entirely new set of neighbors or completely left the droplet. This is indicative of a more liquid-like droplet with a fast exchange rate with the outside environment. Details of this neighbor persistence calculation can be found in the supporting information.

We compared the MSD and neighbor persistence of the target binding proteins across increasing intra-protein attractions *ϵ*_TT_ in Fig 12 and Fig 13, respectively. Not phase separated systems show normal three dimensional diffusion, while proteins in systems that formed condensed droplets have decreased MSD, consistent with transformation from a gas to liquid phase. More interestingly, we see a slow down in the diffusion and longer neighbor persistence time as *ϵ*_TT_ increases from 2.0 to 3.0*k*_B_*T*. Shown in Fig 13A, at *ϵ*_TT_ = 3.0*k*_B_*T* the system with 128 targets appears to be almost solid-like with some neighbors maintained longer than the evaluated time interval. The same decrease in diffusion and increase in neighbor persistence with higher target-target attraction is also seen in the presence of binding polymers as shown in Fig 12B and 13B. Although only non-specific binding polymers are shown in Fig 12B, trends are similar for lock-and-key binding polymers. These results match well with experimental evidence that when RNA-binding proteins are more attracted to themselves, it results in slower protein diffusion times and more solid-like droplets [30]. This also aligns well with evidence that ALS-associated mutations in the FUS protein result in more solid-like, less dynamic droplets as found experimentally in Patel et al. [42] From these results, we expect that any modifications that make these binding proteins more attractive to themselves, such as additional hydrophobic residues in the IDRs of RNA-binding proteins, could lead to solidification of droplets. The increase we see in neighbor persistence could also correspond to lower diffusion-limited reaction rates because neighboring proteins exchange more slowly.

**Fig 12.**
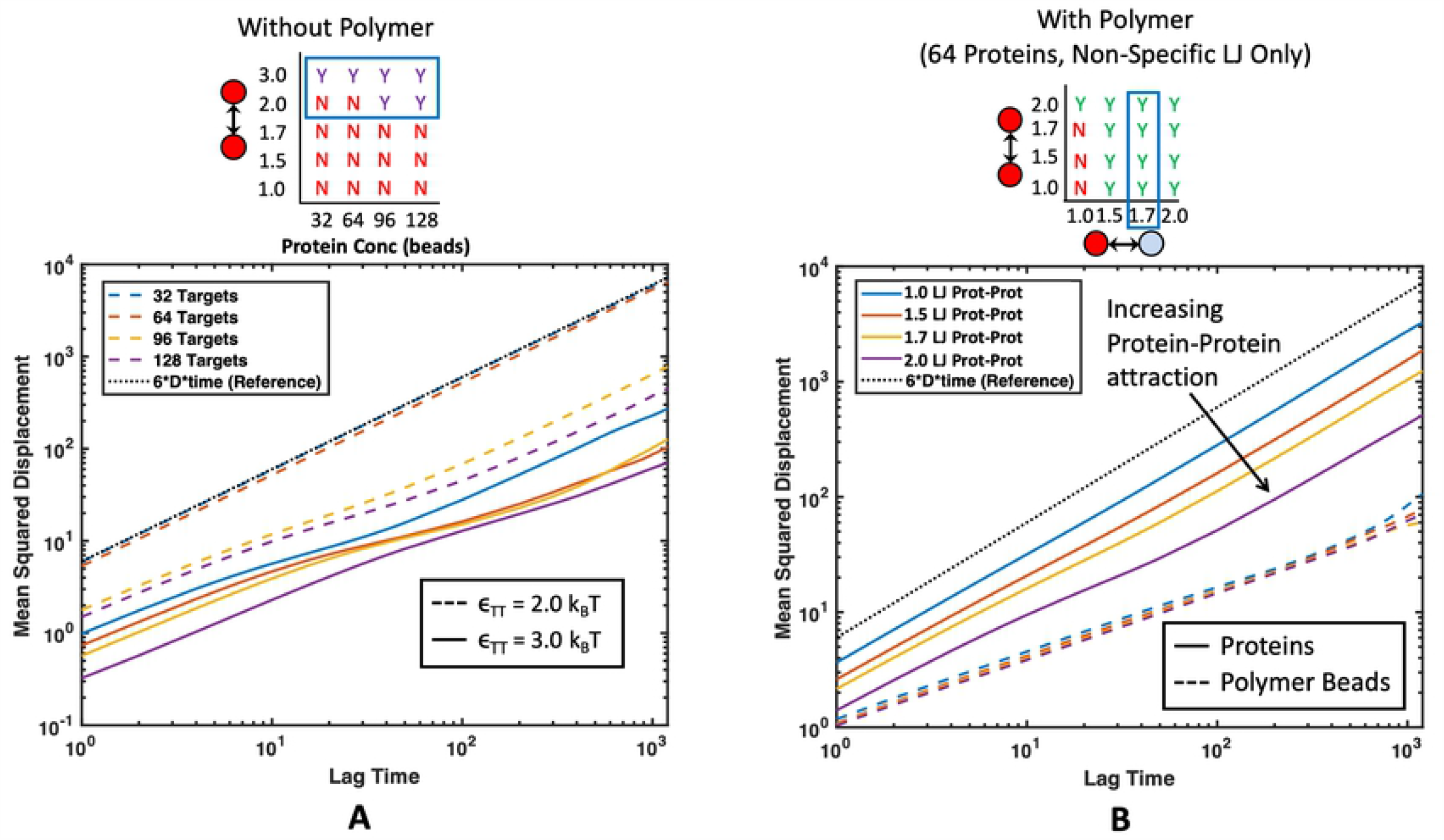
Average mean squared displacement (MSD) of all proteins for several different protein-protein affinities. Corresponding regions of the phase diagrams are highlighted with a blue rectangle above each plot. The black dotted line (------) represents normal 3-D Brownian diffusion. (A) MSD of pure proteins without polymers present. Colors represent different protein concentrations and line pattern represents intra-protein affinity. Not phase separated proteins diffuse with normal Brownian motion whereas phase separated proteins see much slower diffusion rates. Higher *ϵ*_TT_ leads to lower MSD and slower protein diffusion. (B) MSD for 64 proteins interacting with 16mer polymers through non-specific attraction at *ϵ*_TP_ = 1.7. Color corresponds to *ϵ*_TT_. Average MSD for all proteins is shown with a solid line (——) and average MSD for all polymer beads is shown with a dashed line (---). In the presence of polymers, higher intra-protein attraction still leads to slower protein diffusion times.

**Fig 13.**
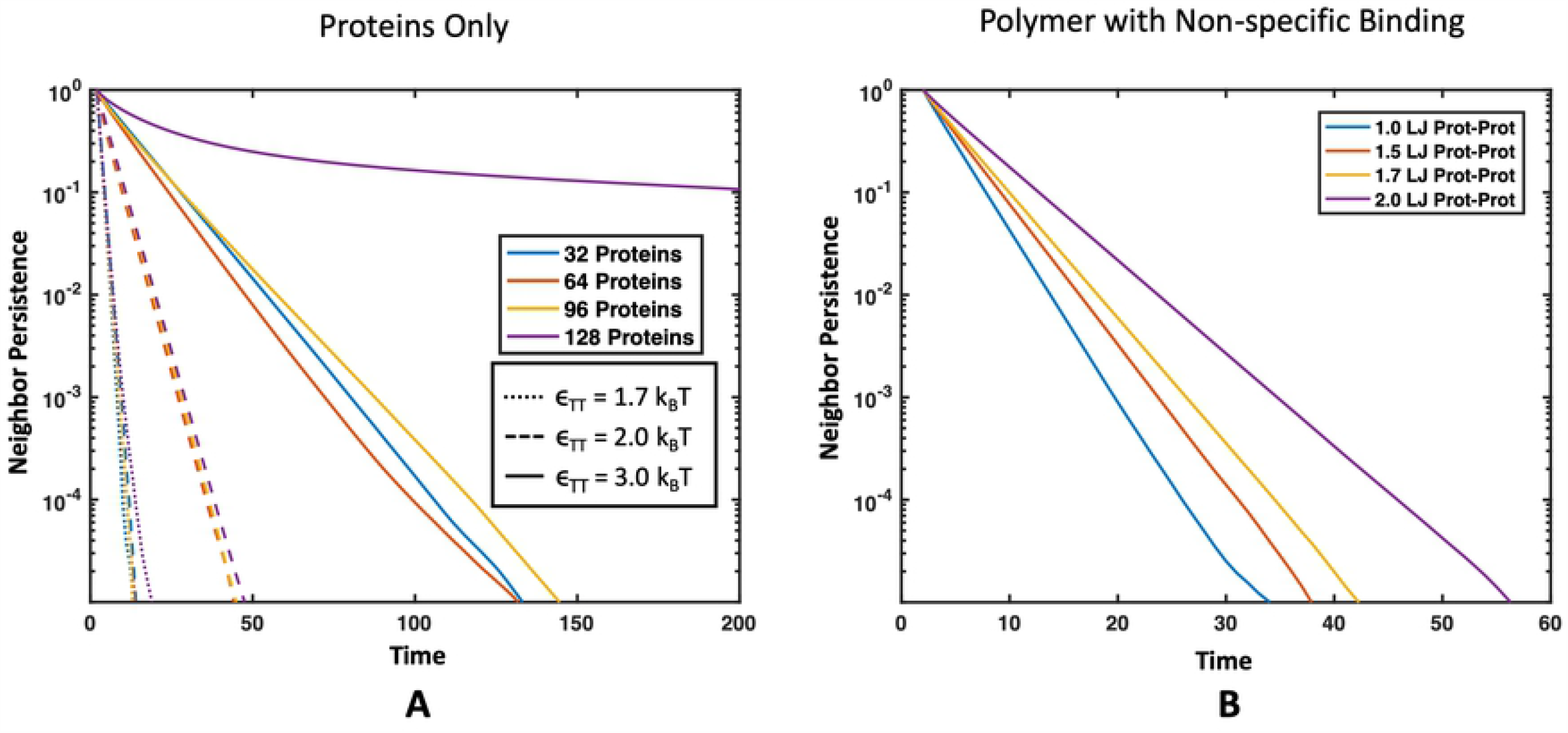
Effect of intra-target protein attraction on neighbor persistence in droplets. Neighbor persistence of a binding protein (A) without a polymer present and with a polymer present that interacts through non-specific interactions. In both cases the time proteins spent with the same neighbors is lengthened as the intra-target attraction *ϵ*_TT_ increases. (A) Line color corresponds to protein concentration and line pattern denotes *ϵ*_TT_. (B) Line color denotes *ϵ*_TT_ in simulations with 64 proteins interacting with four 16mer polymers in theta solvent with *ϵ*_TP_ = 1.7*k*_B_*T*.

Li *et al*. showed experimentally that valency and binding affinity of molecules inside droplets inversely correlate with fluorescence recovery after photobleaching (FRAP) kinetics, which is exactly what we see in simulation [5]. Here, we provide further evidence of this phenomenon through simulation where increasing the affinity of the targets to the polymer decreased the MSD of targets and increased their neighbor persistence. This effect is demonstrated where both non-specific attraction to the polymer and specific valence-limited binding result in a slow down in the MSD in Fig 14 and longer neighbor persistence in Fig 15. The slow down in dynamics caused by higher affinity binding to the polymer is concentration dependent. Unsurprisingly, a higher ratio of binding proteins to polymer results in the polymer having less influence over the droplet dynamics. We also compared the diffusion time of protein beads found within droplets which we considered as any protein with at least 1 neighbor. By comparing the diffusion time and neighbor persistence of these condensed proteins with and without polymers in Fig 16, we found that our simulations also match previous experiments; dynamics of pure protein droplets are slowed upon the addition of a long polymer as shown in experiments with RNA polymers [30]. While we did not see any increase in protein diffusion upon the addition of binding polymers as seen in Maharana et al., this may be because we were not deep enough into the attractive intra-protein energy regime where targets phase separate alone, such as our case where proteins formed a solid-like structure at *ϵ*_TT_ = 3.0*k*_B_*T*. It is possible that this discrepancy is also because the polymers we simulated were relatively long compared with experiments, resulting in slower dynamics for the polymers themselves [15, 30]. Exploring the conditions under which polymers can speed the diffusion of proteins in condensed droplets would be an interesting avenue for future work. In addition, if would be interesting to consider beads of mismatched sizes. Here, we consider beads of equal sizes, but if we used different sizes to disrupt the packing, we might see more significant changes in diffusion.

**Fig 14.**
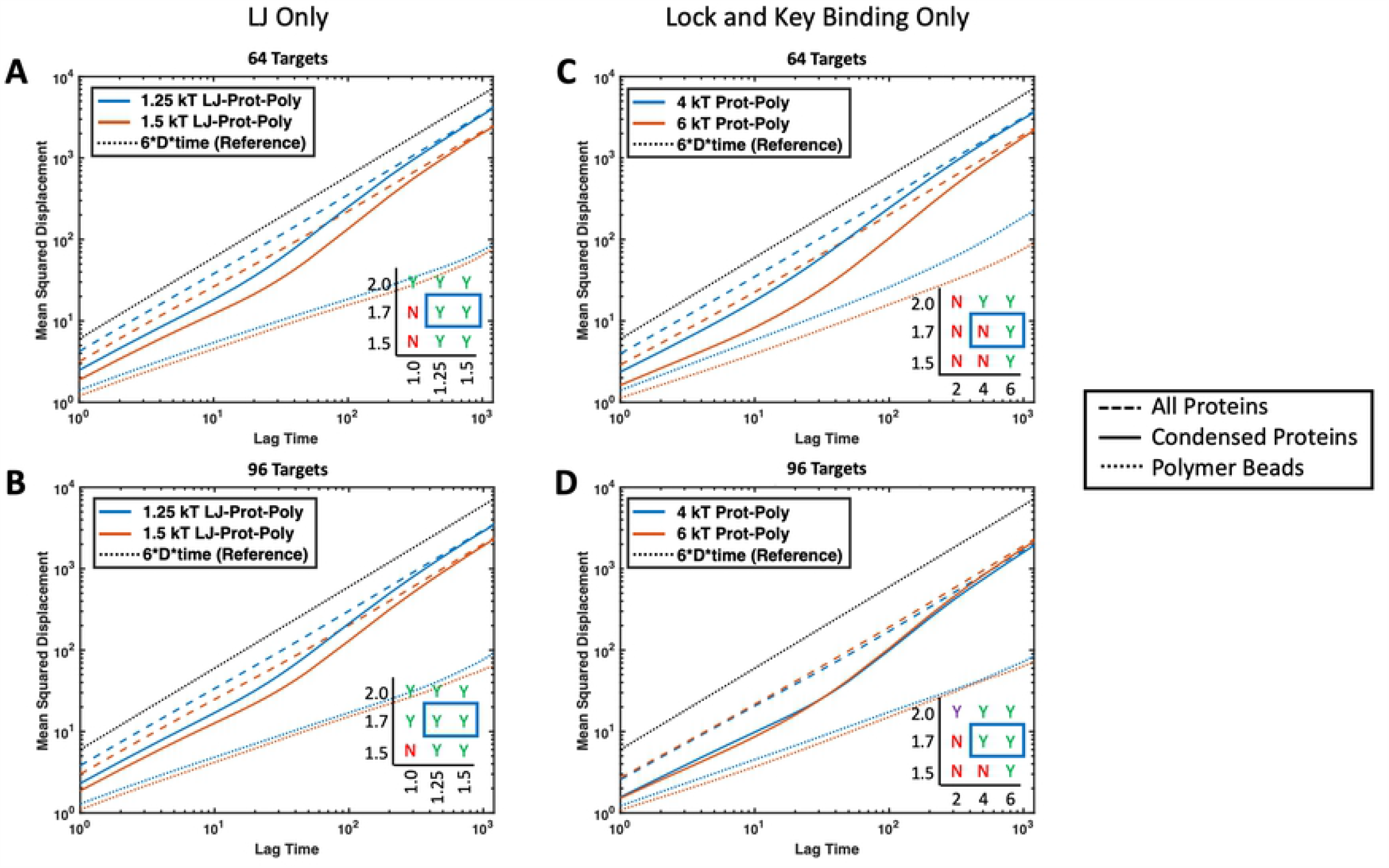
Effect of protein-polymer attraction on MSD of proteins. Average MSD for proteins with *ϵ*_TT_ = 1.7*k*_B_*T* interacting with four 16mer polymers in theta solvent. Dotted black line (------) represents normal Brownian diffusion, dashed lines (---) represent the average MSD over all proteins in the simulation, solid lines (——) represent the average MSD over all proteins that started with at least one neighbor at the beginning of the time interval, and the colored dotted (------) lines represent the average MSD over all polymer beads in the simulation. Colors represent two attraction energies between protein targets and polymers with blue denoting lower affinity than orange. Each plot contains the corresponding phase diagrams with the plotted regions highlighted with a blue rectangle. Cases plotted include (A) non-specific binding polymer with 64 targets and *ϵ*_TP_ = 1.25 and 1.5*k*_B_*T*, (B) non-specific binding polymer with 96 targets and *ϵ*_TP_ = 1.25 and 1.5*k*_B_*T*, (C) specific binding polymer with 64 divalent targets and Δ*E*_0_ = −4 and −6*k*_B_*T*, and (D) specific binding polymer with 96 divalent targets and Δ*E*_0_ = −4 and −6*k*_B_*T*. Protein diffusion slows with increasing protein-polymer attraction, but the polymer has less influence on droplet dynamics when the ratio of proteins to polymer is high.

**Fig 15.**
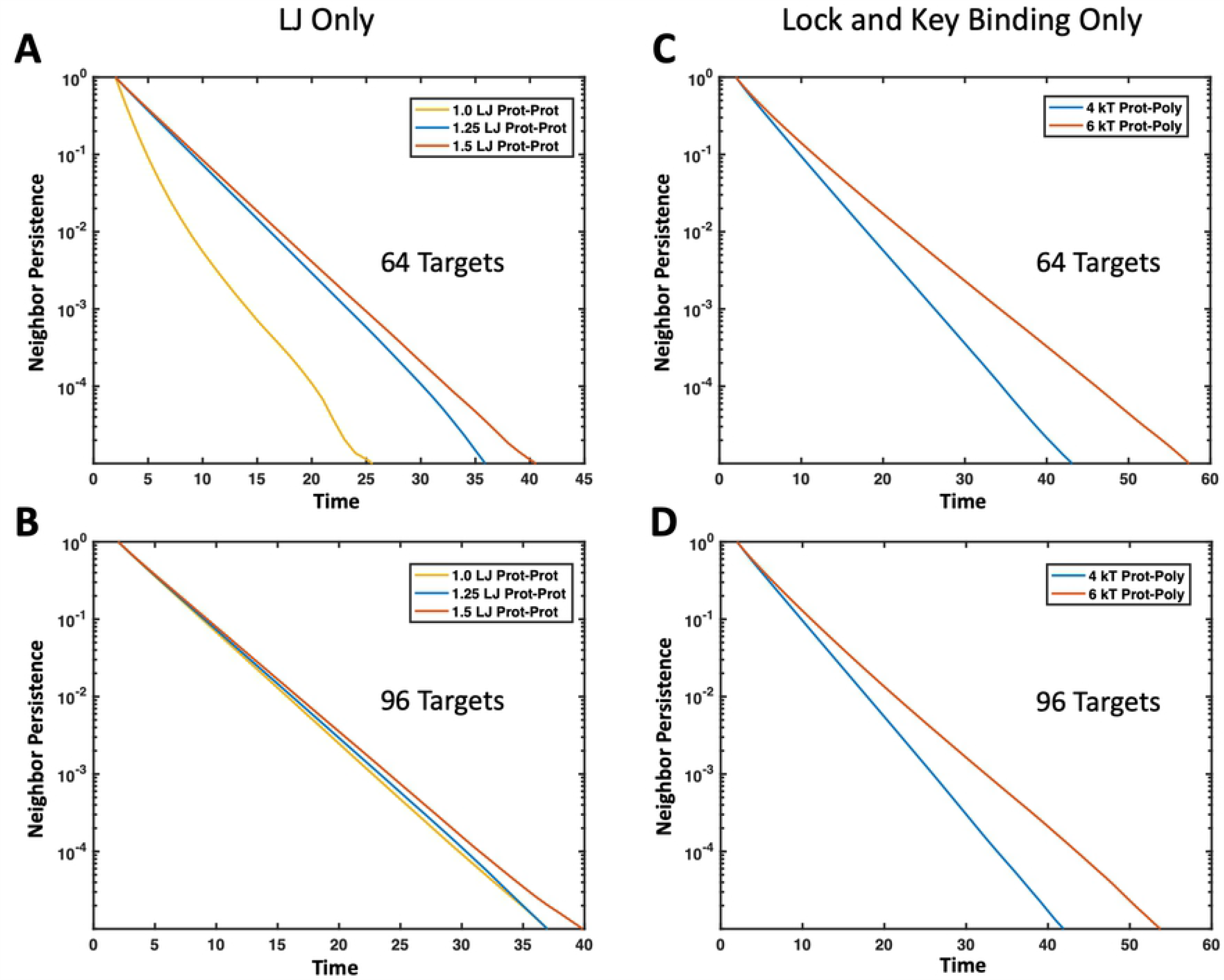
Effect of polymer-target protein attraction on neighbor persistence in droplets. Average time proteins spend with the same neighbors normalized by the average number of initial neighbors. Faster decays to zero indicate a more liquid-like droplet where proteins can move through or exit the droplet freely. Increasing binding affinity to the polymer results in longer protein neighbor persistence. Results are shown for the same cases as Fig 14. (A) Non-specific binding polymer with 64 targets and *ϵ*_TP_ = 1.25 and 1.5*k*_B_*T*, (B) non-specific binding polymer with 96 targets and *ϵ*_TP_ = 1.25 and 1.5*k*_B_*T*, (C) specific binding polymer with 64 divalent targets and Δ*E*_0_ = −4 and −6*k*_B_*T*, and (D) specific binding polymer with 96 divalent targets and Δ*E*_0_ = −4 and −6*k*_B_*T*.

**Fig 16.**
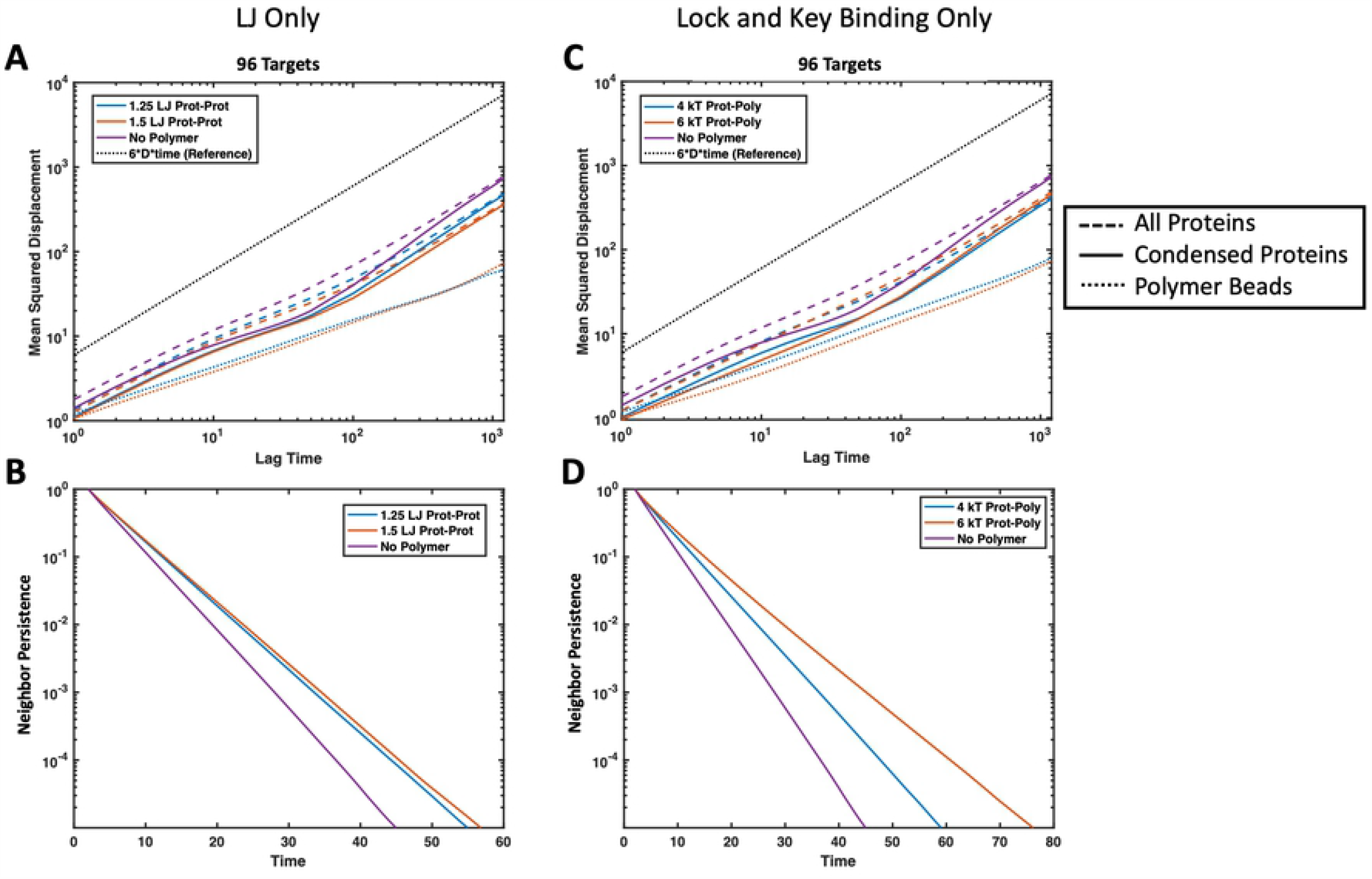
Comparison of protein droplet dynamics with and without polymers. Average MSD (top) and neighbor persistence (bottom) of phase separated system with protein targets binding to four 16mer polymers compared with the dynamics of a phase separated system of pure protein target beads with no polymer (purple). The protein targets interact with *ϵ*_TT_ = 2.0*k*_B_*T*, and all simulations contain 96 target proteins. (A) MSD of proteins experiencing non-specific attraction with the polymers *ϵ*_TP_ = 1.25 (blue) and 1.5*k*_B_*T* (orange). (B) Neighbor persistence of all 96 proteins experiencing non-specific attraction with the polymers *ϵ*_TP_ = 1.25 (blue) and 1.5*k*_B_*T* (orange). (C) MSD of divalent proteins with specific binding to the polymers with Δ*E*_0_ = −4 (blue) and −6*k*_B_*T* (orange). (D) Neighbor persistence of all 96 divalent proteins with specific binding to the polymers with Δ*E*_0_ = 4 (blue) and 6*k*_B_*T* (orange). In MSD plots, the dotted black line (------) is reference for normal Brownian diffusion, dashed lines (---)) represent the average MSD over all proteins in the simulation, solid lines (——) are the average MSD over all proteins that started with at least one neighbor at the beginning of the time interval, and the colored dotted (------) lines represent the average MSD over all polymer beads in the simulation.

## Conclusion

Understanding how multivalent polymers can alter and control the phase transitions of biocondensates and their dynamics is crucial to understanding pathological aggregation.

Here, we present simulations and explore the aggregation and diffusion kinetics of smaller species that bind multivalently to longer polymers. Although our coarse-grain system is relatively simple, we found that we captured the co-phase separation of multivalent polymers and binding proteins found in biocondensates. When comparing to native biocondensates, our system most closely resembles scenarios with two interacting asymmetrically-valent species such as a many-valent polymer binding to a target with low valency. Although there are multiple scenarios that could be represented as our coarse-grain system, one possible example includes RNA as the multivalent polymer and RNA binding proteins (RBPs) such as FUS, hnRNPs, and TDP-43 as the low valency protein target. These RBPs have 1, 2, and 3, RNA recognition motifs (RRMs), and if we imagine the RRMs as specific binding sites, could represent the our monovalent, divalent, and trivalent targets with specific lock-and-key binding sites [18, 36].

Despite the different geometries, our results also seem to align well with systems of two similarly size linear multivalent polymers and their phase transitions [5, 6]. We similarly found that the addition of a multivalently binding polymers can lower the phase boundary for a protein, and that increasing protein valency and binding affinity also lower the phase boundary. Unlike previous research, we also consider the differences in non-specific binding that might come from IDRs versus the specific valency limited binding that come from RRMs in our example biocondensate. We show that changes in the affinity of non-specific interactions can cause more drastic changes in the phase boundary than valence-limited lock-and-key type bonds. Together, they can be used to carefully tune the phase boundary.

Next, we showed that both surface tension and binding affinity could be used to tune droplet order in a system of only three components. When proteins had higher attraction to multivalent polymers, droplets remained mixed with proteins and polymers distributed throughout. When proteins had higher attraction to themselves than to the multivalent polymer, we were able to recreate systems of concentric droplets. Pure proteins formed a central core, while the polymers were pushed into an outer shell. This could have implications for understanding how changes in polymer-protein binding can impact biocondensate function.

Last, we showed that increasing attraction between targets themselves and between targets and polymers can slow the diffusion of targets within condensates and make them more solid-like, consistent with previous experimental results [5, 30]. The attraction to the polymer has a greater effect on target dynamics in droplets at lower target concentrations, when the ratio of polymer beads to targets is higher. After nucleation and growth of a condensed target phase where targets outnumber polymer binding sites, target-target attraction dominates the droplet dynamics. This suggests that, in addition to polymer-binding interactions, changes in the non-specific attraction between binding proteins themselves can induce droplets to be more liquid or more solid-like. Changes in dynamics could have big implications for the reversibility of condensate formation and for reaction rates inside condensates, leading to clear implications for neurodegenerative diseases related to dysregulation of LLPS such as ALS.

While more research needs to be done on specific systems and systems with more than two components we hope that the results presented contribute to the understanding and control of biocondensates and their associated diseases.

## Supporting information

**S1 Fig. Monovalent binding affinity**. Fraction of time a monovalent target spent bound to monomers versus the number of available binding monomers. In the schematic, the target is shown in red and the monomers are shown in light grey-blue. Simulation values are shown as circles (o) with the dashed line (----) showing the fit of the Langmuir adsorption curve for *K*_D_ = 0.1 mM. This figure is adapted from Zumbro *et al*. with permission [39].

**S2 Fig. Example of a typical system energy profile over time**. The total energy is shown for a system with four 16mer polymers and 96 divalent targets with a target-target attraction *ϵ*_TT_ = 1.7*k*_BT_. Simulation energy is shown every 10000 timesteps. There is an initial large drop in energy while the system equilibrates. Production research data is taken from the last quarter of the simulation, past this equilibration time period. This figure is adapted from Zumbro *et al*. with permission from Elsevier [16].

## Acknowledgments

The authors were supported by the Department of Defense (DoD) through the National Defense Science and Engineering Graduate (NDSEG) Fellowship Program. The authors were also supported by the Ida M. Green Fellowship and the Collamore-Rogers Fellowship through the MIT Office of the Dean of Graduate Education.

